# Cbl and Cbl-b Ubiquitin Ligases are Essential for Intestinal Epithelial Stem Cell Maintenance

**DOI:** 10.1101/2023.05.17.541154

**Authors:** Neha Zutshi, Bhopal C. Mohapatra, Pinaki Mondal, Wei An, Benjamin T. Goetz, Shuo Wang, Sicong Li, Matthew D. Storck, David F. Mercer, Adrian R. Black, Sarah P. Thayer, Jennifer D. Black, Chi Lin, Vimla Band, Hamid Band

**Author notes:** Corresponding Authors: Hamid Band and Vimla Band Address: Eppley Institute for Research in Cancer, 986805 Nebraska Medical Center Omaha, NE 68198-6805. Division of Surgical Oncology, LSU Health Sciences and Ochsner LSU Health, Shreveport, LA. Co-first authors (Neha Zutshi, Bhopal C. Mohapatra).

## Abstract

Among the signaling pathways that control the stem cell self-renewal and maintenance vs. acquisition of differentiated cell fates, those mediated by receptor tyrosine kinase (RTK) activation are well established as key players. CBL family ubiquitin ligases are negative regulators of RTKs but their physiological roles in regulating stem cell behaviors are unclear. While hematopoietic *Cbl/Cblb* knockout (KO) leads to a myeloproliferative disease due to expansion and reduced quiescence of hematopoietic stem cells, mammary epithelial KO led to stunted mammary gland development due to mammary stem cell depletion. Here, we examined the impact of inducible *Cbl/Cblb* double-KO (iDKO) selectively in the Lgr5-defined intestinal stem cell (ISC) compartment. *Cbl/Cblb* iDKO led to rapid loss of the *Lgr5*^Hi^ ISC pool with a concomitant transient expansion of the *Lgr5*^Lo^ transit amplifying population. LacZ reporter-based lineage tracing showed increased ISC commitment to differentiation, with propensity towards enterocyte and goblet cell fate at the expense of Paneth cells. Functionally, *Cbl/Cblb* iDKO impaired the recovery from radiation-induced intestinal epithelial injury. *In vitro*, *Cbl/Cblb* iDKO led to inability to maintain intestinal organoids. Single cell RNAseq analysis of organoids revealed Akt-mTOR pathway hyperactivation in iDKO ISCs and progeny cells, and pharmacological inhibition of the Akt-mTOR axis rescued the organoid maintenance and propagation defects. Our results demonstrate a requirement for *Cbl/Cblb* in the maintenance of ISCs by fine tuning the Akt-mTOR axis to balance stem cell maintenance vs. commitment to differentiation.

## INTRODUCTION

Receptor tyrosine kinases (RTKs) and their peptide growth factor ligands function as versatile regulators of stem cell maintenance and cell fate decisions during embryonic development as well as in adult tissues (Mele et al., 2019; Ramachandra et al., 2016; Takahashi et al., 2020). Thus, mechanisms that positively or negatively regulate RTK function have a strong potential to impinge on stem cell homeostasis and fate decisions. Unraveling these regulatory mechanisms at the organismic and molecular levels is imperative to fully understand the biological underpinnings of organ and tissue development, homeostasis and regeneration as well as how disease affects these processes.

Biochemical and cell biological studies have established the Cbl (<u>C</u>asitas <u>B</u>-Lineage <u>L</u>ymphoma) family of ubiquitin ligases (Cbl, Cbl-b and Cbl-c), as activation-dependent negative regulators of RTKs and non-receptor tyrosine kinases linked to cell surface receptors (Mohapatra et al., 2013). Cbl proteins possess canonical tyrosine kinase-binding (TKB) and RING finger domains separated by a conserved linker, regions required for their ubiquitin ligase activity, with Cbl and Cbl-b including an extended C-terminal region with proline-rich sequences, tyrosine phosphorylation sites and a ubiquitin-associated domain, a region lacking in Cbl-c. Furthermore, Cbl and Cbl-b are expressed in overlapping patterns across many tissues while Cbl-c mRNA expression is primarily localized to epithelial tissues (Mohapatra et al., 2013; Griffiths et al., 2003). While all three Cbl-family proteins function as negative regulators of tyrosine kinases *in vitro*, only Cbl and Cbl-b have emerged as key regulators of physiological functions in vivo (Mohapatra et al., 2013). *Cblc*-null mice show no overt phenotypes, and a protein product of the *Cblc* gene has not been categorically identified (Griffiths et al., 2003). *Cbl-*null mice show impairment of spermatogenesis and thymic development along with mildly hypercellular hematopoietic organs (Murphy et al., 1998; Naramura et al., 1998; El Chami et al., 2005), and exhibit a lean phenotype due to hyperactive insulin receptor signaling (Molero et al., 2004). *Cblb-*null mice are developmentally normal, but their immune cells are hyper-responsive to antigenic stimulation thus evoking autoimmunity (Chiang et al., 2000; Bachmaier et al. 2000). *Cblb*-null mice are also resistant to disuse muscle atrophy due to hyperactive IGF1R signaling (Nakao et al., 2009). Thus, Cbl and Cbl-b are clearly physiologically-relevant as negative regulators of RTK signaling.

While *Cbl* or *Cblb* null mice have a relatively normal lifespan under pathogen-free laboratory conditions, combined germline *Cbl* and *Cblb* gene deletion is embryonic-lethal, supporting their redundant roles during embryonic development. This redundancy is further demonstrated by a rapidly-lethal inflammatory disease upon combined *Cbl* and *Cblb* deletion in T-cells (Naramura et al., 2002 and Goetz et al., 2016) and a lethal myeloproliferative disease upon combined deletion in hematopoietic stem cells (HSC) (Naramura et al., 2010; An et al., 2015; An et al., 2016). These latter studies in HSCs showed that Cbl proteins are required to regulate physiological levels of HSC proliferation, and their deletion led to reduced HSC quiescence and impaired ability to reconstitute hematopoiesis upon bone marrow transplantation (An et al., 2015). *In vitro* studies using shRNA depletion have shown a role for Cbl and Cbl-b in maintaining asymmetric neural stem cell division (Ferron et al., 2010). Recently, we observed that mammary epithelium-specific *Cbl/Cblb* double deletion led to substantial retardation of the postnatal mammary gland development, with depletion of mammary stem cells (MaSCs) (Mohapatra et al., 2017). Contrary to the impact of *Cbl/Cblb* double deletion in HSCs, no apparent hyper-proliferation of intermediate progenitors or accumulation of progeny cells was observed in the double knockout (DKO) mammary glands. Furthermore, the fate of *Cbl/Cblb*-null MaSCs remained unclear primarily due to lack of clear-cut systems to track the stem-progenitor-mature cell hierarchy in the mammary gland (Mohapatra et al., 2017). Given the well-established stem-progeny cell relationships in the intestinal epithelium, we have undertaken a comprehensive examination of the role of *Cbl* and *Cblb* in regulating ISC homeostasis.

The intestinal epithelial cell monolayer is a major environment-organism interface and is responsible for critical functions of digestion and absorption of food while maintaining a barrier against microbes. The intestinal epithelium is organized to ensure its maintenance throughout adult life in the face of rapid turnover of the mature, functional epithelial cells that line the villi in the small intestine. It is estimated that the entire epithelium is renewed every 3-5 days (Barker et al., 2014). The homeostatic maintenance of such a rapidly-renewing tissue requires a well-regulated program to rapidly generate mature epithelial cell types from intestinal stem cells (ISCs) located at the base of the crypts. These ISCs divide daily to form rapidly-cycling progenitor cells to expand the number of progeny cells. The progenitors further migrate along the crypt-villus axis as they adopt specific differentiated fates of secretory, absorptive, or other specialized cell types. Thus, the intestinal epithelial stem cell program provides an elegant system in which to explore mechanisms that control adult epithelial stem cell maintenance in the context of epithelia, an area of great interest since epithelial malignancies account for most of human cancers (Andersson-Rolf et al., 2017).

Recent advances have elucidated the general hierarchy of ISCs and how they help maintain the epithelium under homeostasis or following extrinsic insults that lead to rapid loss of epithelial integrity or that eliminate rapidly-dividing stem and progenitor cell pools (Takeda et al., 2011; van Es et al., 2012; Buczaki et al., 2013; Metcalfe et al., 2014). It is now well-accepted that crypt-base columnar cells (CBCs) are rapidly-dividing stem cells that maintain the epithelium under homeostatic conditions (Barker et al., 2007). A less abundant and relatively slowly-dividing ISC population located at the +4 position with respect to crypt base is thought to serve as a reserve pool that is rapidly recruited to replenish the more actively-cycling CBC-ISCs when the latter are eliminated, such as upon gamma radiation exposure (Takeda et al., 2011; van Es et al., 2012; Buczaki et al., 2013). Additional studies suggest that the slowly-dividing ISCs, commonly identified as label-retaining cells, may differentiate into secretory cells under homeostatic conditions (Buczaki et al., 2013). Other studies point to the plasticity of these populations, with interconversion between the rapidly-dividing and the slowly-cycling cells (Takeda et al., 2011; van Es et al., 2012; Buczaki et al., 2013). There is considerable overlap in the markers that have been used to define the various ISC populations, but the WNT target gene product Lgr5 (Leucine Rich Repeat-Containing G Protein-Coupled Receptor 5) is generally accepted as a marker of the majority of long-lived but rapidly-cycling ISCs (Barker et al., 2007; Munoz et al., 2012). Lgr5 is a multi-pass cell surface receptor for the R-spondin family of growth factors that enhances WNT signaling, a pathway required for ISC proliferation and colorectal oncogenesis (Barker et al., 2009; Carmon et al., 2011; 2012). Aside from WNT signaling, other signaling pathways, including those mediated by Notch, BMP, and growth factor receptor tyrosine kinases (RTKs), are known to function as key regulators of ISCs and intestinal epithelial maintenance (Clevers 2013; Barker and Tan, 2015).

Substantial evidence supports the prominent roles of RTKs and/or their ligands in intestinal development, maintenance, and disease. Newborn epidermal growth factor receptor (EGFR)-null mice, on a genetic background that allows live pups to be born, display gross defects in the intestinal mucosa along with reduced cell proliferation (Miettinen et al., 1995; Sibilia & Wagner, 1995). The ISC niche role of the Paneth cell requires secretion of EGFR ligands by these cells (Sato et al., 2011). Application of EGF to intestinal mucosa of whole animals enhances epithelial cell proliferation (Cheung et al., 2009). The importance of EGFR-mediated signals in maintaining a proliferating pool of ISCs is further established by recent studies of intestinal epithelial organoid cultures in which inhibition of EGFR activation induced the Lgr5+ ISCs to enter quiescence (Basak et al., 2017). RTKs of the EphB family, fibroblast growth factor receptor (FGFR), and insulin-like growth factor receptors (IGFRs) have also been shown to play crucial roles during gut morphogenesis, mucosal proliferation, crypt survival, and differentiation and positioning of cells along the crypt-villus axis (Batlle et al., 2002; Holmberg et al., 2006; Vidrich et al., 2009; Al Alam et al., 2015; Freier et al., 2005). *Lrig1*, a negative regulator of the ErbB family and other RTKs (Powell et al., 2012; Wong et al., 2012), was shown to be enriched within the crypt compartment, and *Lrig1*-null mice display hyper-proliferation of intestinal epithelium and duodenal adenomas as early as 3 months of age (Powell et al., 2012). On the other hand, loss of Shp2, a positive regulator of RTK signaling, was shown to cause expansion of Paneth cells along with an increase in the Lgr5+ stem cells due to an imbalance of MAPK and Wnt pathways (Heuberger et al., 2014). Additionally, altered differentiation and loss of stemness have been reported upon overexpression of FGF10 (Al Alam et al., 2015). These studies support the potentially dichotomous roles of RTK signaling in the maintenance of intestinal epithelial homeostasis.

Currently, nothing is known about the physiological roles of Cbl-family proteins in the regulation of ISCs. Previous studies have suggested that Lrig1 may function in part through its interaction with Cbl to promote RTK ubiquitination and degradation (Gur et al., 2004). Thus, the dramatic intestinal epithelial hyperproliferation and duodenal adenoma formation phenotypes in *Lrig1*-null mice (Powell et al., 2012) supported the possibility that Cbl and Cbl-b may be involved in the regulation of ISCs. Here, we use *Cbl* and *Cblb* gene deletion in the Lgr5+ ISC compartment to provide evidence that Cbl and Cbl-b are redundantly required to maintain Lgr5+ cycling ISCs and that Cbl/Cbl-b deletion leads to rapid loss of ISCs through enhanced Akt-mTOR-dependent differentiation of ISCs into progeny cells followed by unequal lineage differentiation. Our studies therefore reveal a novel and previously unanticipated requirement for a tyrosine kinase negative regulatory mechanism provided by Cbl and Cbl-b in the maintenance of cycling ISCs.

## RESULTS

### CBL and CBL-B expression is enriched in intestinal crypts compared to villi

Since the roles of Cbl-family proteins in the intestine have not been examined previously, we first assessed the expression pattern of Cbl and Cbl-b in the intestine (Fig. 1A-E). Immuno-histochemical (IHC) staining of formalin-fixed and paraffin-embedded intestinal tissue sections from 6-week old wildtype (WT) C57Bl/6 mice revealed higher Cbl expression in crypts with lower signal in the villus area; lack of any crypt and villus epithelium staining of the Cbl*^-/-^* mouse intestinal sections established the specificity of Cbl staining (Fig. 1A). IHC for Cbl-b showed a similar expression pattern, although the relative difference between the crypt and villus signals was less pronounced (Fig. 1B). To confirm these findings, we analyzed lysates of crypt and villus fractions of intestinal mucosal tissue by western blotting (WB) (Fig.1C). As expected, the levels of phospho-histone3 (pH3) and Cyclin D1, markers of crypt-associated cell proliferation (Van Landeghem et al., 2015; Shinozaki et al., 2003), were higher in the crypt fraction whereas Villin-1 (El Marjou et al., 2004; Wang Y et al., 2008) was enriched in the villus fraction (Fig.1C). Notably, WB confirmed the crypt vs. villus enrichment of Cbl and Cbl-b. The crypt-enrichment of Cbl and Cbl-b expression is reminiscent of that of the RTK negative regulator Lrig1 previously shown to partially function through Cbl (Gur et al., 2004; Wong et al., 2012; Powell et al., 2012).

**Fig. 1.**
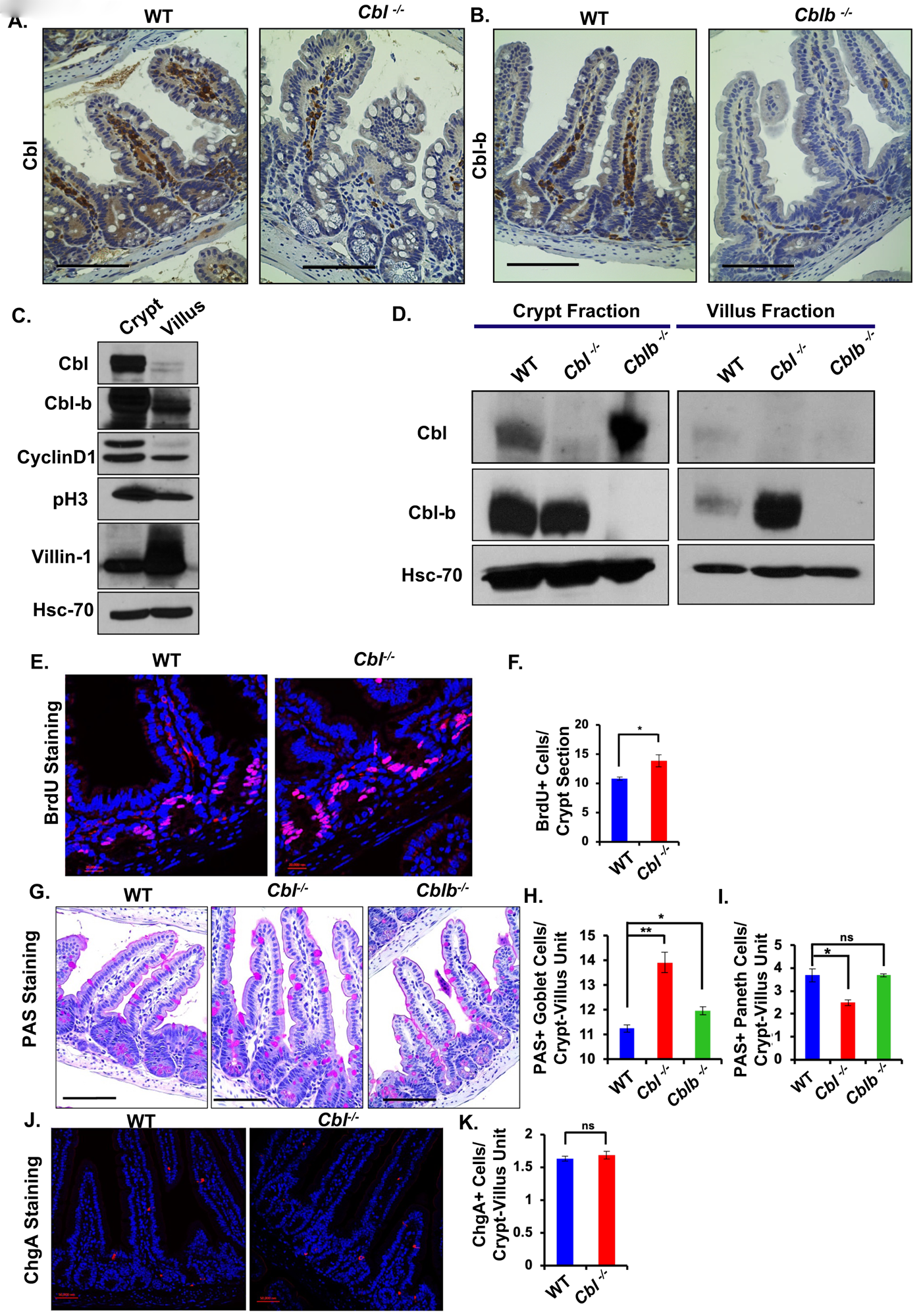
Preferential expression of Cbl and Cbl-b proteins in intestinal crypts areas of the intestine. (A, B) Immunohistochemical (IHC) staining for Cbl (A) and Cbl-b (B) on formalin-fixed paraffin-embedded (FFPE) tissue sections from 6-weeks old mice reveals their expression in a gradient from crypt to villus in WT animals (left panels); lack of epithelial staining in sections from *Cbl^-/-^* or *Cblb^-/-^* knockout mice (right panels) demonstrates the specificity of antibody staining. (C) Crypt-villus fractionation of intestinal mucosa followed by immunoblotting confirmed the crypt enrichment of Cbl and Cbl-b expression. Blotting for crypt (pH3 and Cyclin D1) and villus (Villin-1) associated markers confirmed the purity of villus and crypt fractions. Hsc-70 served as a loading control. (D) Immunoblot analysis of crypt and villus fractions prepared from age- and gender-matched *WT*, *Cbl^-/-^*and *Cblb*^-/-^ animals. Within the crypt compartment, deletion of *Cblb* resulted in an increase in the expression of Cbl while Cbl-b expression was unchanged upon deletion of *Cbl*. The reverse was seen in the villus compartment, with an increase in Cbl-b expression upon *Cbl* deletion whereas Cbl expression was unchanged in *Cblb*^-/-^ animals. Scale bars in A and B are 100 µm; data are representative of 3 experiments. Altered intestinal epithelial cell proliferation and differentiation in *Cbl^-/-^* mice. (E) *WT* and *Cbl^-/-^* mice were injected with BrdU 4 hours prior to euthanasia and FFPE sections of intestines were stained with anti-BrdU antibody (Red) and DAPI (Blue); scale bar= 20 mm. (F) Quantification of the number of BrdU+ cells per crypt shows an increased proportion of cells in S phase of the cell cycle in Cbl null mice. (G-I) PAS staining of tissue sections from age matched *WT*, *Cbl^-/-^* and *Cblb^-/-^* mice revealed a significant increase in the number of goblet cells (PAS+ cells in the villus) in both *Cbl*^-/-^ and *Cblb*^-/-^ mice (H) and a modest reduction in the Paneth cell (PAS+ cells in the crypt) number in *Cbl*^-/-^ mice (I), scale bar= 100 µm. (J, K) Chromogranin A staining (Red), showed no significant difference in the number of enteroendocrine cells between WT and *Cbl^-/-^*mice; scale bar= 50 mm. Data are represented as +/- SEM of 4 independent experiments. Statistical analysis used *student’s two-tailed t-test*; ns, p ≤ 0.05, *; p ≤ 0.01, **.

WB of crypt and villus fractions of WT vs. *Cbl^-/-^* or *Cblb^-/-^* single knockout mice revealed comparable Cbl-b levels in WT and *Cbl^-/-^* crypts, but a marked elevation of Cbl levels in *Cblb^-/-^* crypt fractions as well as an increase in Cbl-b levels in *Cbl^-/-^* the villi fractions (Fig. 1D). The overlapping crypt-enriched Cbl and Cbl-b expression and compensatory changes in their expression upon deletion of the other gene supported the likelihood of redundant Cbl/Cbl-b roles in the intestinal epithelium, as we observed during hematopoiesis (Naramura et al., 2010; An et al., 2015). Due to lack of authenticated antibodies against Cbl-c, we examined the Cbl-c expression indirectly using the *Cblc^-/-^* mice with a *LacZ* reporter driven by *Cblc* promoter (Griffiths et al., 2003). Co-staining with X-gal (substrate for β-galactosidase) and Ki67 (to demarcate the crypt compartment) showed that *Cblc* (*LacZ*) was predominantly localized to villi and non-overlapping with the Ki67+ crypts (Supplementary Fig. S1A). Given the predominant Cbl/Cbl-b expression in crypts and Cbl-c expression in differentiated villus cells, our further studies focused on the roles of Cbl and Cbl-b in the context of ISCs.

### Altered intestinal epithelial cell proliferation and differentiation in *Cbl^-/-^* mice

Given the intestinal crypt-enriched expression of Cbl and Cbl-b (Fig.1A-D), we first examined the status of cell proliferation and differentiation in the intestinal epithelium of 6-week-old *Cbl^-/-^* or *Cblb^-/-^*mice compared to age-matched WT controls. Assessment of bromo-deoxyuridine (BrdU) incorporation demonstrated an increase in cell proliferation in the *Cbl^-/-^*mouse intestinal crypts compared to WT mice, suggesting increased stem/progenitor cell proliferation (Fig. 1E, F). On the other hand, immunofluorescence staining for pH3 revealed no significant difference in proliferating crypt epithelial cells between *Cblb^-/-^* and WT mice (Supplementary Fig. S1B, C). Hematoxylin and Eosin (H&E)-stained sections demonstrated an intact overall intestinal epithelial architecture in *Cbl^-/-^* and *Cblb^-/-^* mice, but with increased abundance of goblet cells (Supplementary Fig. S1D). PAS staining to directly visualize goblet and Paneth cells (van Es et al., 2012) revealed a moderate or mild goblet cell hyperplasia in *Cbl^-/-^*or *Cblb^-/-^* mouse intestines, respectively, (Fig. 1G, H), and a reduction in Paneth cell numbers in *Cbl^-/-^* compared to WT mice (Fig. 1G, I). Chromogranin A staining for neuroendocrine cells (Gunawardene et al., 2011) showed no significant differences between *Cbl^-/-^* and WT mice (Fig. 1J, K). Collectively, the increased crypt epithelial cell proliferation in *Cbl^-/-^*mice and the altered numbers of goblet in *Cbl^-/-^* and *Cblb^-/-^*mice and Paneth cells in *Cbl^-/-^*mice suggested a functionally significant role of Cbl and Cbl-b play in ISC proliferation and differentiation, with a more prominent role for Cbl.

### Inducible Lgr5+ ISC-specific *Cbl* knockout on a *Cblb^-/-^* background leads to expansion of progenitors with shrinkage of the stem cell compartment

Given the results presented above, we generated an ISC-specific gene deletion model to directly examine the role of Cbl and Cbl-b in the Lgr5+ ISCs, which are required for homeostatic maintenance of the intestinal epithelium and known to self-renew while cycling. To circumvent the embryonic lethality of constitutive DKO of *Cbl* and *Cblb* in mice (Naramura et al., 2002), we crossed the *Cbl^flox/flox^*; *Cblb^-/-^*mice previously used for *Cbl/Cblb* DKO in HSCs (Naramura et al., 2010; An et al., 2015; An et al. 2016) or MaSCs (Mohapatra et al., 2017) with mice harboring the *Lgr5-EGFP-IRES-creERT2* and *R26-LSL-LacZ* reporter knock-in alleles (Barker et al., 2007) to generate the *Cbl^flox/flox^*; *Cblb^-/-^*; *Lgr5-EGFP-IRES-creERT2*; *Rosa26-LacZ* mice. In these mice, Lgr5+ ISC-specific inducible DKO of *Cbl* and *Cblb* (iDKO) can be achieved upon tamoxifen-induced *Cre* activation and deletion of floxed-*Cbl* alleles, and lineage tracing of double KO ISCs can be monitored by X-Gal staining. Lgr5 promoter-driven GFP expression also allowed us to track the fate of Lgr5+ ISCs based on the levels of their GFP expression.

Following three consecutive daily intraperitoneal injections of TAM, mouse intestines were analyzed on day 5 or 10 (Fig. 2A). Quantitative real-time PCR (Q-PCR) of isolated GFP+ intestinal epithelial cell RNA confirmed the *floxed-Cbl* deletion at both time-points (Supplementary Fig. S2A; Fig. 2B-C). Consistent with the specificity of *Lgr5-cre* expression (Barker et al., 2007), loss of *Cbl* expression was seen in GFP-high (GFP Hi) *Lgr5*+ stem cells and their GFP-low (GFP Lo) daughter cells but not in GFP-negative (GFP neg) cells, the latter representing the non-ISC population and ISCs in which *Lgr5-cre* was not expressed due to well-established mosaic *Lgr5-cre* expression (Barker et al., 2007). To assess the impact of iDKO on Lgr5+ ISC maintenance, we quantified the GFP Hi (stem) and GFP Lo (progenitor) intestinal epithelial cell populations in Control vs. iDKO mice by flow cytometry 5- or 10-days after initiating TAM induction (Fig. 2D-G; gating strategy outlined in Supplementary Fig. S2A). The iDKO mice exhibited a significant reduction in the percentage of GFP Hi ISCs and a concomitant increase in the percentage of GFP Lo progenitor cells at both 5- and 10-day time points (Fig. 2E, G). A similar trend was seen when absolute cell numbers were quantified (Supplementary Fig. S2 B-C). Q-PCR for stemness-associated genes *Lgr5* and *Olfm4* confirmed the gating strategy and demonstrated reduced *Lgr5* and *Olfm4* expression in the GFP-Hi ISC compartment of iDKO compared to control mice (Supplementary Fig. S2 D-E). These results suggested that concurrent *Cbl* and *Cblb* deletion leads to a rapid shrinkage of the Lgr5+ stem cell compartment, with a concomitant expansion of the transit-amplifying compartment.

**Fig. 2.**
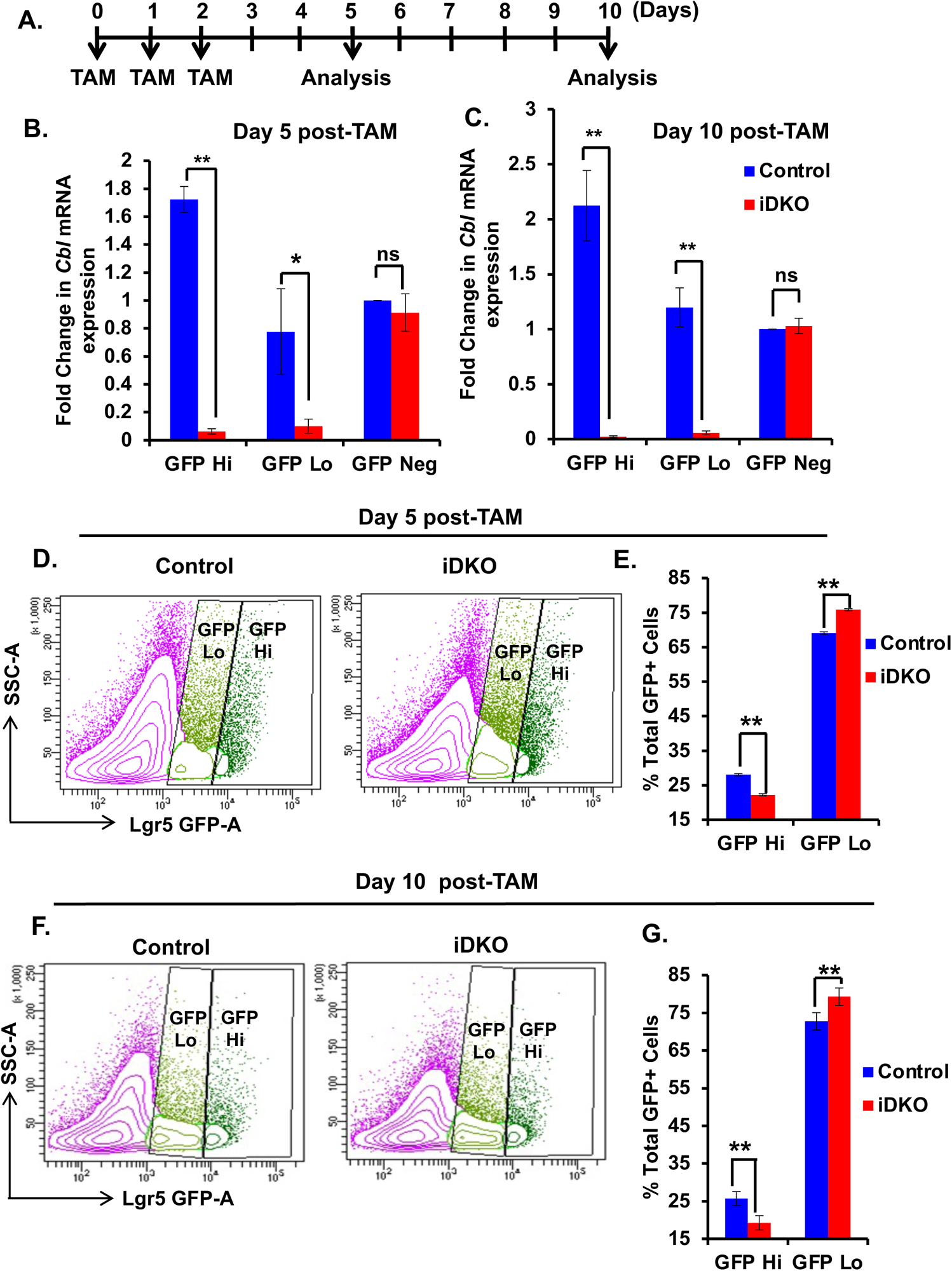
Expansion of the intestinal epithelial progenitor compartment at the expense of stem cells upon inducible *Cbl/Cblb* double KO in *Lgr5+* stem cells. (A) Schematic showing the experimental plan followed. *Cbl^flox/flox^*; *Cblb^-/-^*; *Lgr5-cre^ERT2^*; *R26-LacZ* (iDKO) mice received tamoxifen (TAM) on days 0, 1 and 2 to induce conditional deletion of floxed *Cbl* alleles in *Lgr5+* cells on a *Cblb* null background. (B-C) Deletion of *Cbl* was assessed using real time PCR of FACS-sorted GFP-high and GFP-low vs. GFP-negative epithelial cells at day 5 and day 10 of induction. (D-G) Flow cytometric analysis of live epithelial cells showed a significant expansion of *Lgr5-GFP* Lo (progenitor) cells and a concomitant decrease in *Lgr5-GFP* Hi (stem cells) at 5 days (D-E) and 10 days (F-G) after TAM induction in *Cbl/Cblb* iDKO vs. Control mice. Data are represented as mean +/- SEM of 4 independent experiments. Statistical analysis using *student’s two-tailed t-test* is shown; ns, p ≤ 0.05, *; p ≤ 0.01, **.

### *Cbl/Cblb* iDKO mice exhibit increased cell proliferation in intestinal crypts

As we observed a significant depletion of Lgr5+ stem cells upon *Cbl/Cblb* iDKO, we examined the status of crypt cell proliferation in Control vs. iDKO mice by quantifying BrdU-incorporating cells specifically in GFP+ crypts, representing those in which *Lgr5-cre* was active (Barker et al., 2007). The iDKO GFP+ crypts showed a significant increase in BrdU+, proliferating cells compared to those in control animals at day 5 of TAM induction (Fig. 3A, B), with a similar albeit less pronounced trend at day 10 (Fig. 3C, D). Immunostaining for cleaved caspase 3 did not show any differences in the frequency of apoptotic cells in GFP+ crypts of iDKO vs. control mice (Fig. 3E-H); a higher relative number of apoptotic cells in both genotypes at day 5 relative to day 10 of TAM treatment is consistent with the reported transient toxicity of TAM on proliferating cells (Zhu et al., 2013). Thus, concurrent loss of Cbl and Cbl-b in Lgr5+ ISCs is associated with increased cell proliferation.

**Fig. 3.**
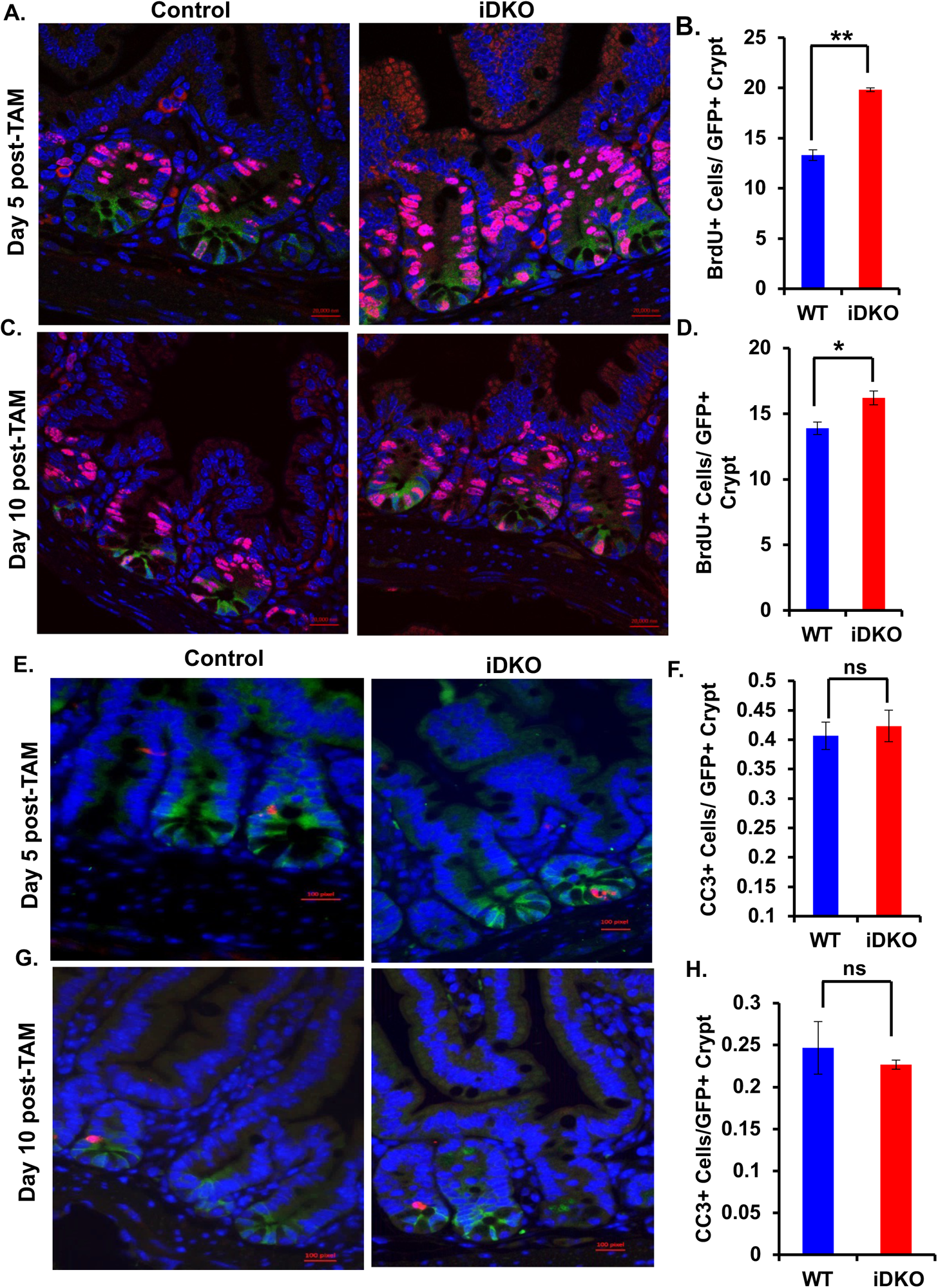
Increased crypt cell proliferation with no change in apoptosis upon Lgr5+ cell-specific *Cbl/Cbl-b* iDKO in the intestine. (A-D) Control and *Cbl/Cblb* iDKO mice were injected with BrdU 4 hours prior to euthanasia and FFPE sections of intestines were stained for BrdU (Red), GFP (Green) and DAPI (Blue). Quantification of BrdU+ cells within GFP+ crypts showed a significant increase in the fraction of iDKO stem/ progenitor cells in S phase of the cell cycle at day 5 (A-B) and day 10 (C-D) of initiating TAM treatment. (E-H) Cleaved caspase 3 (Red) co-immunostaining with GFP (Green) and DAPI (Blue) revealed no significant difference in the number of dead cells between control and iDKO mice at day 5 (E-F) or day 10 (G-H) after initiating TAM treatment. Analyses at each time point were repeated three times. Data are presented as mean +/- SEM with statistical analysis using *student’s two-tailed t-test*; ns, p ≤ 0.05, *; p ≤ 0.01, **. Scale bar= 20 mm.

### Loss of Cbl*/* Cbl-b promotes commitment to differentiation

As the progenitor cells arising from Lgr5+ ISCs are known to proliferate faster (Barker et al., 2007), and we observed an expansion of the GFP Lo (Lgr5-low) progenitors with a reduction in the GFP Hi (Lgr5+) stem cells upon iDKO (Fig. 2), the increased crypt cell proliferation without any change in apoptosis (Fig. 3) suggested an increased commitment of the iDKO Lgr5+ ISCs into differentiated progeny. To assess if this is the case, we performed lineage tracing of the Lgr5+ stem cells in iDKO and Control mice by tracking their *LacZ+* progeny after TAM induction (Barker et al., 2007). At day 5 and 10 time points, we observed more *LacZ+* cells in the crypts and villi with a significant increase in the percentage of partially or completely blue crypts in iDKO compared to control mice (Fig. 4A-D); this trend persisted at day 10 after induction, albeit only the increase in the percentage of fully blue crypts remained significant (Fig. 4C, D). Notably, the number of crypts with a single blue cell was significantly reduced in iDKO mice at day 10 of TAM induction (Fig. 4D); such single *LacZ+* cells within crypts likely represent quiescent Lgr5+ cells, possibly those that become secretory progenitors without further division (Buczaki et al., 2013; Takeda et al., 2011). The quiescent progenitor pool is known to express high levels of *Dll1* (Delta like ligand 1) and *Ngn3* (Neurogenin 3) (van Es et al., 2012; Schonhoff et al., 2004). Q-PCR analysis revealed a marked alteration of these markers within the sub-fractions of Lgr5+ cells in iDKO mice, with an elevated *Dll1* and *Ngn3* expression in the GFP Hi (Lgr5-Hi ISCs) fraction and a decrease in their expression in the GFP Lo (Lgr5-low progenitor) fraction (Fig. 4E-F).

**Fig. 4.**
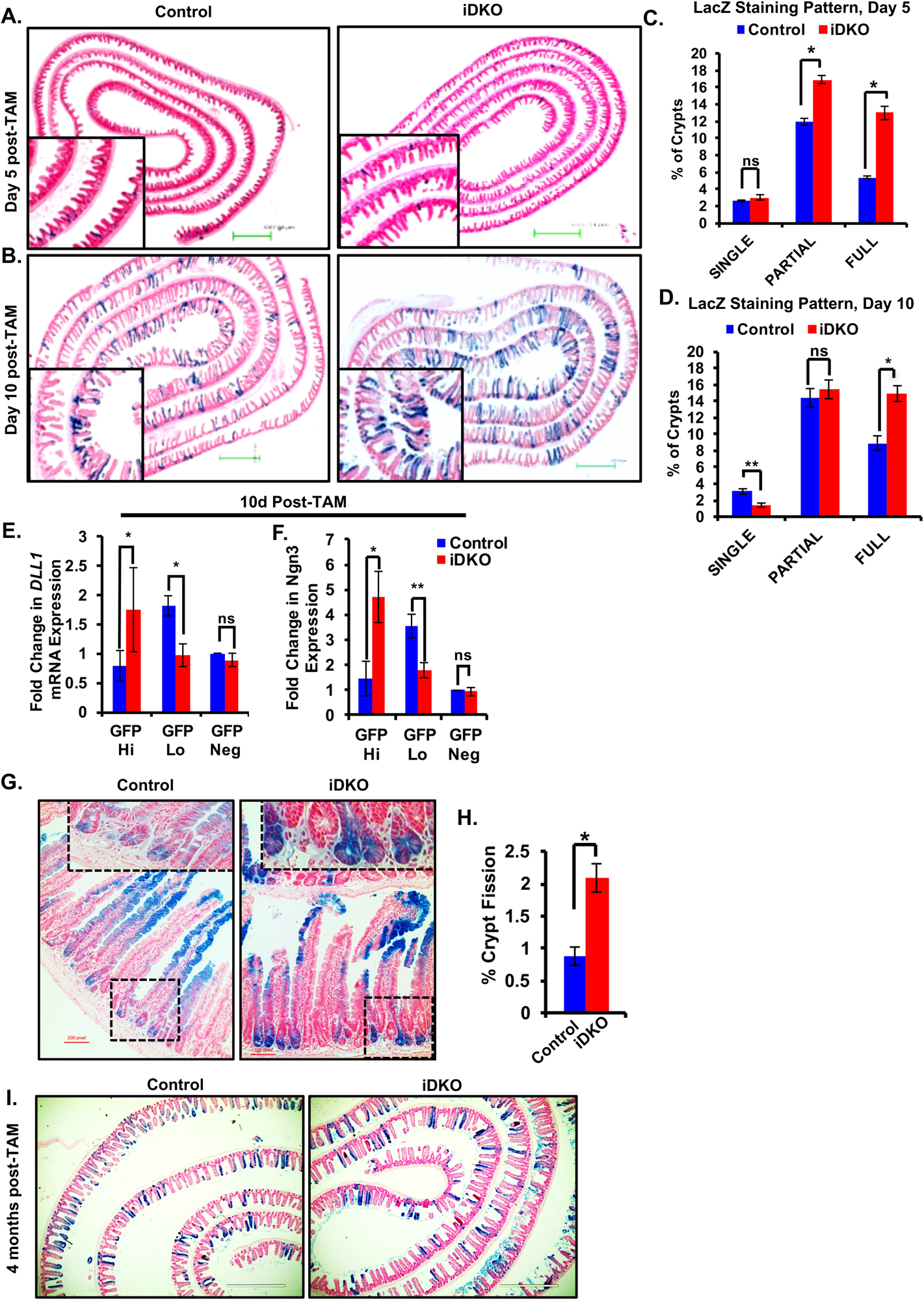
Increased commitment to differentiation upon Lgr5+ cell-specific *Cbl/Cbl-b* iDKO in the intestine. (A-B) X-Gal staining of intestinal sections at 5 (A) and 10 (B) days post-TAM induction to trace the fate of Lgr5+ ISCs shows an increase in blue progeny in *Cbl/Cblb* iDKO mice at both time points as compared to control mice; scale bar= 1,000 µm. (C-D) Quantitative analysis of the staining shown in A-B was performed by counting 2,000 crypts per genotype and assessing the number of crypts with a single blue cell or with fully or partially blue crypts. An increase in the percentage of full and partial blue crypts was observed together with a reduction in the percentage of crypts with single blue cells. (E-F) Real-time PCR analysis shows a mis-localization of the quiescent secretory progenitor markers Delta like 1 (*Dll1*) (E) and Neurogenin 3 (*Ngn3*) (F) in GFP Hi and GFP Lo populations in *Cbl/Cblb* iDKO vs. control mice. (G-H) Enumeration of the number of LacZ+ crypts undergoing fission shows an increase in iDKO mice at 10 days post-TAM induction as compared to control mice. (I) LacZ staining at 4 months post-TAM induction showed comparable staining between control and *Cbl/Cblb* iDKO mice with no signs of hyperplasia in the latter; scale bar= 400 µm. Analyses at each time point were repeated four times. Data are presented as mean +/- SEM with statistics using *student’s two-tailed t-test*. ns, p ≤ 0.05, *; p ≤ 0.01, **.

Many *LacZ+* crypts in iDKO mice appeared as clusters, suggesting increased crypt fission. Enumerating the % of *LacZ+* crypts undergoing fission indeed revealed this to be the case in the iDKO mice (Fig. 4G, H). Notably, analysis at 4 months of TAM induction revealed no differences in LacZ staining of crypts in iDKO vs. WT mice, and we observed normal architecture throughout the gastrointestinal tract with no signs of hyperplasia (Fig. 4I), in sharp contrast to adenomas observed in Lrig1-null mouse intestine (Powell et al., 2012).

Given the increased *Dll1* expression in the Lgr5 Hi ISC population, we further assessed the secretory cell fate of the iDKO ISC progeny by combined PAS X-gal staining. As with *Cbl-* or *Cblb* single knockout mice (Fig. 2), the iDKO mice showed a significant increase in the number of LacZ+/PAS+ goblet cells and a concomitant reduction in the number of LacZ+/PAS+ Paneth cells compared to control mice (Supplementary Fig. S3A, B). LacZ and chromogranin A co-staining revealed no significant difference in the number of enteroendocrine cells in iDKO vs. control mice (Supplementary Fig. S3C, D).

Overall, these results support the conclusion that concurrent loss of Cbl and Cbl-b proteins commits the Lgr5+ ISCs and their rapidly dividing progeny towards differentiation, with skewing of the secretory cell fate towards goblet cells.

### *Cbl/Cblb* iDKO mice exhibit delayed recovery from radiation-induced intestinal mucosal injury

Rapid replacement of intestinal epithelial cells shed/lost under normal homeostasis or following mucosal insults, such as exposure to radiation, is critical to maintain an intact epithelial barrier. Radiation treatment induces a rapid loss of cycling Lgr5+ stem and progenitor compartments in the gut with regeneration by re-entry of the quiescent Lgr5+ secretory progenitors into the cell cycle and their rapid expansion to re-establish epithelial homeostasis (Buczaki et al., 2013; van Es et al., 2012). Thus, we reasoned that the impact of the premature differentiation of rapidly dividing Lgr5+ ISCs and their progeny in iDKO mice would be more discernible in a post-radiation recovery setting despite mosaic gene deletion directed by the *Lgr5-cre* (Barker et al., 2007). To avoid concurrent bone marrow dysfunction from whole-body irradiation commonly used to induce the intestinal mucosal injury (Hua et al., 2012), we used focused beam abdominal radiation, known to reduce the hematopoietic toxicity and to effectively target the gut epithelium with 100% recovery through re-establishment of the intestinal epithelium by the quiescent pool of gut stem cells in normal mice (Van Landeghem et al., 2012).

Groups of control and iDKO mice were subjected to 14Gy of focused beam abdominal radiation and TAM treatment as indicated (Fig. 5A) and weighed daily to monitor weight loss, which provides an accurate surrogate indicator of induced mucosal injury in this model (Van Landeghem et al., 2012). While Control mice began to recover their weight by day 5 post-radiation and reached their pre-radiation weight by day 9 (data not shown), iDKO mice lagged significantly (Supplementary Fig. S4 A). H&E staining of intestinal sections harvested on day 7 showed blunted villi and several crypt-less regions in iDKO mice compared to substantially better-preserved histology in control mice (Fig. 5B). Ki67 staining revealed a significant reduction in the number of surviving crypts (defined as a crypt with at least 5 adjacent Ki67+ cells by Metcalf et al., 2014) in iDKO as compared to control mice (Supplementary Fig. S4B, C). FACS analysis at day 5 after radiation demonstrated a reduction in the percentage as well as absolute number of Lgr5-Hi cells and a mild increment in the percentage and number of Lgr5-Lo cells in iDKO compared to control mice (Fig. 5C-E; Supplementary Fig. S4D, E).

**Fig. 5.**
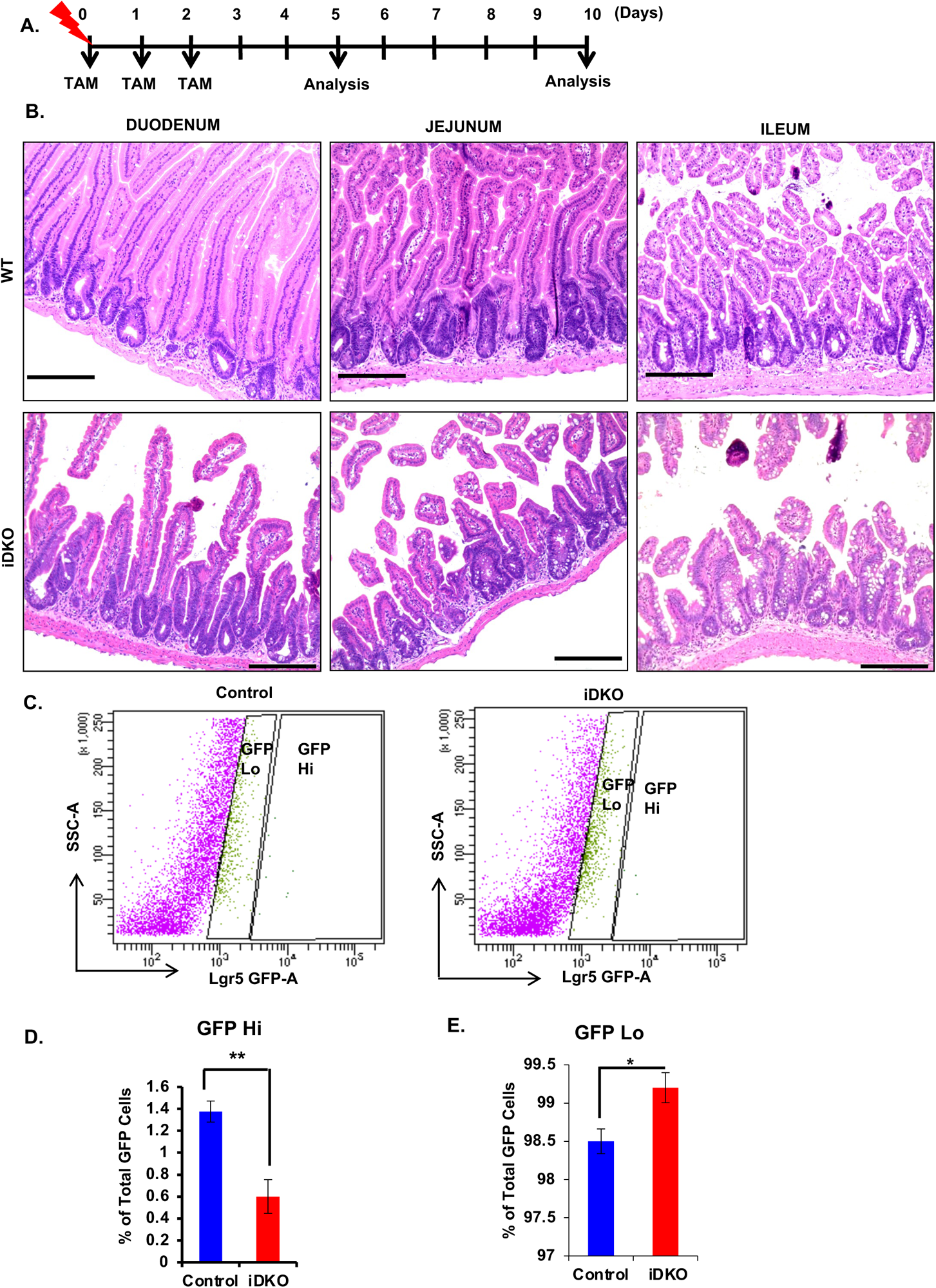
*Cbl/Cbl-b* iDKO mice exhibit delayed recovery from abdominal radiation injury. Control and *Cbl/Cblb* iDKO mice were exposed to CT-based Conformal X-ray radiation (14Gy) focused on the abdomen to assess the ability of intestinal stem/progenitor cells to re-establish epithelial homeostasis post-damage. (A) The schematic shows the experimental plan followed. (B) Histological analysis of H&E-stained sections of different regions of intestine on day 7 post-injury showed hyperproliferating crypts with fission and repopulation of villi in the control mice while *Cbl/Cblb* iDKO mice showed bare, crypt-less patches and blunted villi with little recovery; representative pictures are shown; scale bar = 200 µm. (C-E) The inability of *Cbl/Cblb* iDKO mice to regenerate injured intestinal epithelium is associated with a precipitous drop in the Lgr5-Hi stem cell population (C, D) and a mild increase in the Lgr5-Lo progenitors (C, E) analyzed at d5 post-injury. Analysis at each time point was repeated three times. Data are presented as mean +/- SEM with statistics using *student’s two-tailed t-test;* ns, p ≤ 0.05, *; p ≤ 0.01, **.

### *Cbl/Cblb* iDKO impairs crypt stem cell self-renewal by hyperactivating the Akt-mTOR pathway

Organoid cultures initiated from isolated intestinal epithelial crypts or purified Lgr5+ stem cells provide a facile system to dissect mechanisms of ISC self-renewal (Sato et al., 2009, Sato et al., 2011). Given the mosaic expression of Lgr5-Cre, we examined the crypt epithelium-derived organoids isolated from *Cbl^f/f^; Cblb^f/f^* (double-floxed or FF Control) or *Cbl^f/f^; Cblb^f/f^;* R26-CreERT+ (FF/creERT) mice (Mohapatra et al 2017) and cultured in the presence of 4-hydroxy tamoxifen (4-OH-TAM) for the indicated time points. The FF Control organoids showed a marked increase in crypt (seen as evaginations) and villus domain formation over the 72-h observation period and H&E staining at the end showed a uniform epithelial layer surrounding the more amorphous interior (Fig. 6A, B), as expected (Sato et al., 2009; Miura S et al., 2017). In contrast, the FF/creERT organoids cultured in 4-OH-TAM showed an increase in crypt domain formation at 24 and 48 h time points, but these receded rapidly by 72 h, and H&E staining showed marked disintegration with no clear epithelial layer (Fig. 6A, B). Quantification of the crypt domains confirmed these results (Fig. 6C). Upon replating, the control TAM-treated organoids re-created the original organoid architecture with intact crypt and villus domains; in contrast, no intact organoid structures could be seen upon re-plating of TAM-treated FF/creERT organoids (Fig. 6D). The expected Cbl and Cbl-b deletion was confirmed upon 48 h and 72 h of 4-OH-TAM treatment of FF/creERT organoids (Fig. 6E). Thus, Cbl and Cbl-b are required for intestinal epithelial homeostasis *in vitro*.

**Fig. 6.**
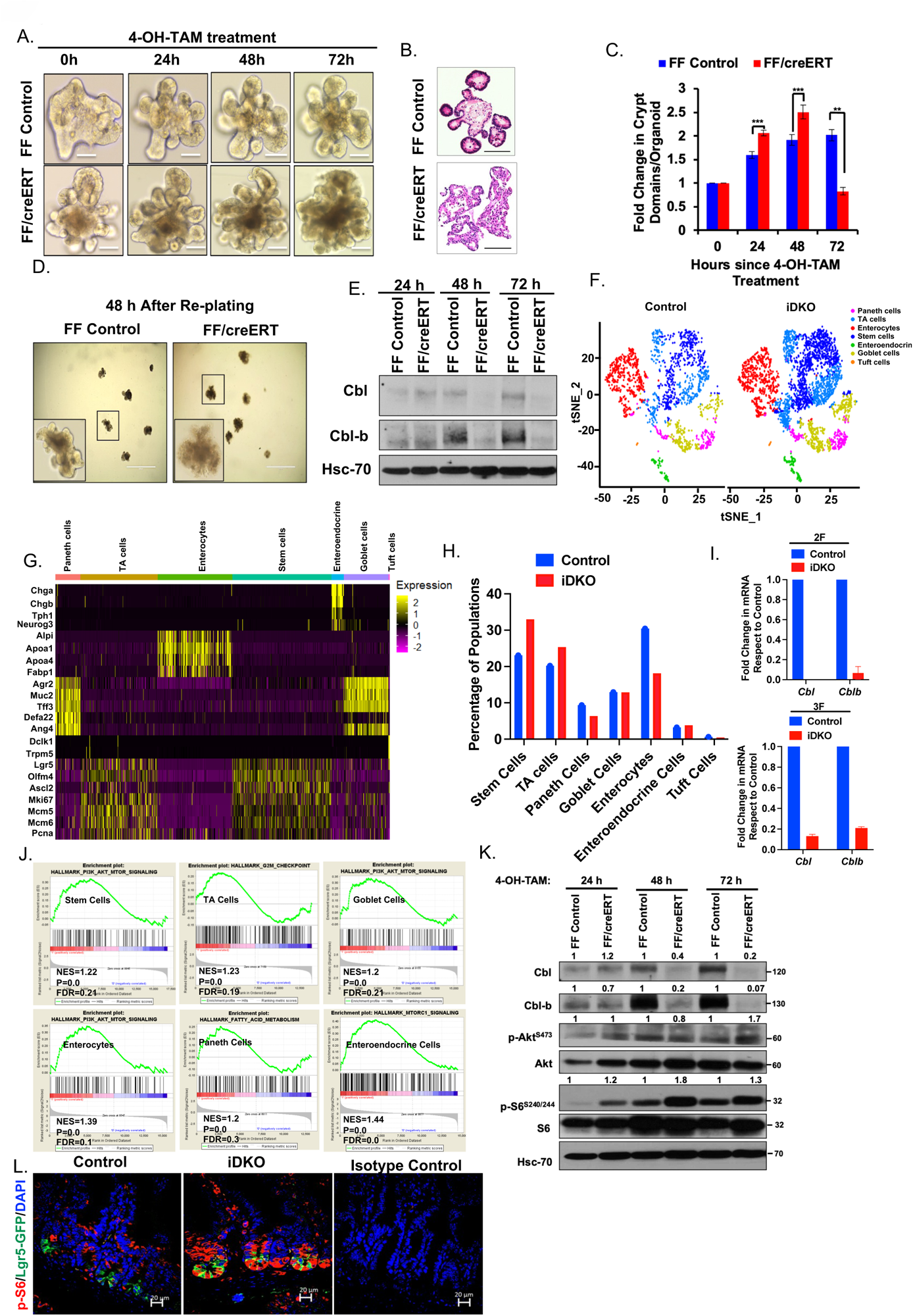
Conditional *Cbl/Cbl-b* deletion impairs self-renewal in intestinal crypt organoid cultures by upregulating the Akt-mTOR pathway. Equal number of crypts isolated from *Cbl*^flox/flox^; *Cblb*^flox/flox^ (FF Control) and *Cbl^flox/flox^*; *Cblb^flox/flox^*; *R26Cre^ERT2^* (FF/creERT) mice were plated in 100% Matrigel in the presence of growth factors to form organoids. Once formed, organoids were re-plated at a 1:4 split ratio and treatment with 400 nM 4-OH TAM was initiated after 24 h to induce *Cbl/Cblb* deletion. (A) Bright field imaging showed that while FF control mouse intestinal organoids exhibited a steady increase in budding (crypt domains) over time (up to 72 hours of observation), the FF/creERT mouse organoids showed increased budding until 48 h after 4-OH-TAM induction but rapidly lost crypt domains by 72 h post-induction; scale bar = 40 µm. (B) Representative H&E-stained images of FF control and FF/CreERT organoids confirm the loss of morphological features in the latter; scale bar = 50 µm. (C) Ten distinct organoids per genotype were followed up to 72 hours and change in the number of buds (crypt domains) at each time point relative to time 0 was quantified. (D) FF Control and FF/creERT organoids grown in 4-OH-TAM for 72 h were re-passaged and imaged after 48 h. Note the lack of organoid structures with intact morphology in FF/creERT organoid cultures, supporting a loss of self-renewal contrary to growth and intact morphology of control organoids; scale bar = 400 µm. (E) Immunoblotting of organoid lysates at different time points confirmed effective *Cbl/Cblb* deletion by 48 and 72 h time points in FF/creERT organoids. HSC70, loading control. FF/creERT organoids derived from two independent female mice (2F and 3F) were cultured in the presence (iDKO) or absence (control) of 400 nM 4-hydroxy-TAM for 72 hours. Single cells isolated from the organoids were processed through the GemCode Single Cell Platform (10X Genomics) to perform single cell RNAseq and analyzed using Seurat. (F) tSNE map of combined controls and combined *Cbl/Cblb* iDKO mouse organoid cells. Cells are grouped into seven clusters based on transcriptome profiles and are colored accordingly. The cell types were assigned based on gene expression profile of stem cells (*Lgr5, Ascl2, Axin2, Olfm4, Gkn3*), transit-amplifying (TA) cells (*Mki67, Cdk4, Mcm5, Mcm6, Pcna*), enterocytes (*Alpi, Apoa1, Apoa4, Fabp1*), Paneth cells (*Lyz1, Defa17, Defa22, Defa24, Ang4*), enteroendocrine cells (*Chga, Chgb, Tac1, Tph1, Neurog3*), Goblet cells (*Muc2, Tff3, Agr2*) and Tuft cells (*Dclk1, Trpm5, Gfi1b*). (G) Heat map shows the expression levels of top cluster-specific genes in each cluster. Yellow represents the highest expression while purple represents low or no expression. (H) Histogram depicting the percentage of cells in each cluster. (I) Validation of *Cbl* and *Cblb* deletion in the 2F and 3F organoids by qPCR. (J) Gene set enrichment analysis (GSEA) of each cell type between control and iDKO organoids shows upregulation the of PI3K-Akt-mTOR signaling pathway upon *Cbl/Cblb* iDKO in stem cells, goblet cells, enteroendocrine cells and enterocytes. NES (normalized enrichment score), FDR (false discovery rate) and p values are indicated in the GSEA plots. (K) To validate the alterations in PI3K-mTOR signaling upon *Cbl/Cblb* iDKO, organoid lysates were collected at different time-points of culture in the presence of 4-OH-TAM and analyzed by immunoblotting. Clear upregulation of p-AKT/mTOR pathway components (p-Akt and p-S6) is seen. Hsc-70 served as a loading control. Densitometries of Cbl, Cbl-b, p-Akt/Akt and p-S6/S6 after normalizing to loading control in comparison to FF control sample in each time point are indicated on the top of the immunoblots. (L) Intestinal tissue sections from Control and *Cbl/Cblb* iDKO mice 10 d after initiating TAM treatment were stained with antibodies against p-S6 to assess the impact of *Cbl/Cblb* deletion. Sections stained with secondary antibody alone were used as a negative control. p-S6 levels (red in color) were sharply increased in iDKO as compared to control sections; scale bar= 20 µm. Quantified data are presented as mean +/- SEM of three independent experiments with statistics using *student’s two-tailed t-test*. ns, not significant; p ≤ 0.05, *; p ≤ 0.01, **; p≤ 0.001, ***.

To gain unbiased mechanistic insights into the nature of molecular alterations caused by *Cbl/Cblb* iDKO, we carried out single cell RNA-seq analysis of cells isolated from untreated (Control) vs. 4-OH-T-treated (*Cbl/Cblb* iDKO) organoid cultures from FF/CreERT mice. Expression of cell type-specific index genes in the transcriptomic profiles identified seven cell clusters corresponding to stem cells (*Lgr5, Ascl2, Axin2, Olfm4, Gkn3*), transit-amplifying (TA) cells (*Mki67, Cdk4, Mcm5, Mcm6, Pcna*), enterocytes (*Alpi, Apoa1, Apoa4, Fabp1*), Paneth cells (*Lyz1, Defa17, Defa22, Defa24, Ang4*), enteroendocrine cells (*Chga, Chgb, Tac1, Tph1, Neurog3*), Goblet cells (*Muc2, Tff3, Agr2*) and Tuft cells (*Dclk1, Trpm5, Gfi1b*) in both control and iDKO organoids (Fig. 6F-H), consistent with the major cell types observed in previous single cell RNA-seq analyses of intestinal epithelial organoids (Grun et al., 2015; Haber et al., 2017). Q-PCR analyses verified the loss of *Cbl* and *Cblb* gene expression in iDKO relative to control organoids (Fig. 6I). Gene set enrichment analyses identified an upregulation of the PI3-kinase/AKT/mTOR pathway in multiple cell compartments in iDKO organoids, including stem cells, goblet cells, enterocytes, and enteroendocrine cells (Fig. 6J). WB analysis of FF Control vs. FF/CreERT organoid lysates confirmed a nearly complete loss of Cbl and Cbl-b expression by 72 hours after adding TAM (Fig. 6K). Immunoblotting showed an upregulation of p-Akt levels in FF/CreERT organoids compared to controls at 24 and 48h that remained sustained at 72h of TAM treatment, and a similar pattern was seen with p-S6 (Fig. 6K). Immunostaining of tissue sections from TAM-induced control vs *Cbl/Cblb* iDKO mice showed that p-S6 levels were markedly increased in iDKO crypts compared to crypts in control mice (Fig. 6L). Collectively, these results supported the conclusion that the AKT/mTOR pathway is hyperactivated upon *Cbl/Cblb* iDKO in ISCs, consistent with the Akt-mTOR pathway hyperactivation upon Cbl/Cbl-b deletion in other systems (Li et al., 2009; Li et al., 2011; Mohapatra et al., 2017; An et al., 2015) and the importance of the the Akt/mTOR pathway downstream of receptor tyrosine kinases in intestinal epithelial homeostasis.

To assess the role of the hyperactive AKT/mTOR pathway in the observed defective intestinal epithelial maintenance upon *Cbl/Cblb* iDKO, we treated FF/creERT organoids with 4-OH-TAM in the absence or presence of specific inhibitors of AKT (MK-2206; 250 nM) and/ or mTOR (Rapamycin; 10 nM). The dual fluorescent reporter of Cre activation (Red before and green after Cre activation) (Muzumdar et al., 2007) in the organoids revealed uniform activation of Cre upon 4-OH-TAM treatment (Fig. 7A). Compared to 4-OH-TAM treated organoids without inhibitors, which showed the expected disruption of the organoid structure and reduced crypt domain number by day 3, MK-2206 treatment led to a small but statistically insignificant preservation of the crypt domains and epithelial integrity (Fig. 7A-B). Notably, treatment with Rapamycin or Rapamycin plus MK-2206 led to a marked rescue of the organoid disruption induced by *Cbl/Cblb* deletion (Fig. 7A-B). Of note, treatment with either inhibitor led to some reduction in crypt domain formation at the 24 h time point (Fig. 7B); however, this effect was independent of *Cbl/Cblb* deletion and also seen with control organoids (Supplementary Fig. S5A-B). Re-plating of FF/creERT organoids cultured under the more effective inhibitor conditions for 72 hours showed a substantial retention of organoid structures when analyzed 48 h after replating (Fig. 7C). GFP reporter expression ruled out the organoids seen after re-plating arising from cells in which Cre activation had not occurred. Western blot analysis revealed the expected upregulation of p-Akt and p-S6 in FF/creERT organoids cultured with 4-OH-TAM alone (Fig. 7D). The levels of p-Akt and p-S6 phosphorylation were drastically reduced in organoids treated with Rapamycin or Rapamycin plus MK-2206 (Fig. 7D). Collectively, the sustained *in vitro* and *in vivo* Akt/mTOR pathway activation, together with the reversal of the disruption of *Cbl/Cblb* iDKO organoid architecture upon Akt/mTOR pathway inhibition support the conclusion that negative regulation of the Akt-mTOR pathway by Cbl and Cbl-b is required to maintain the ISCs.

**Fig. 7.**
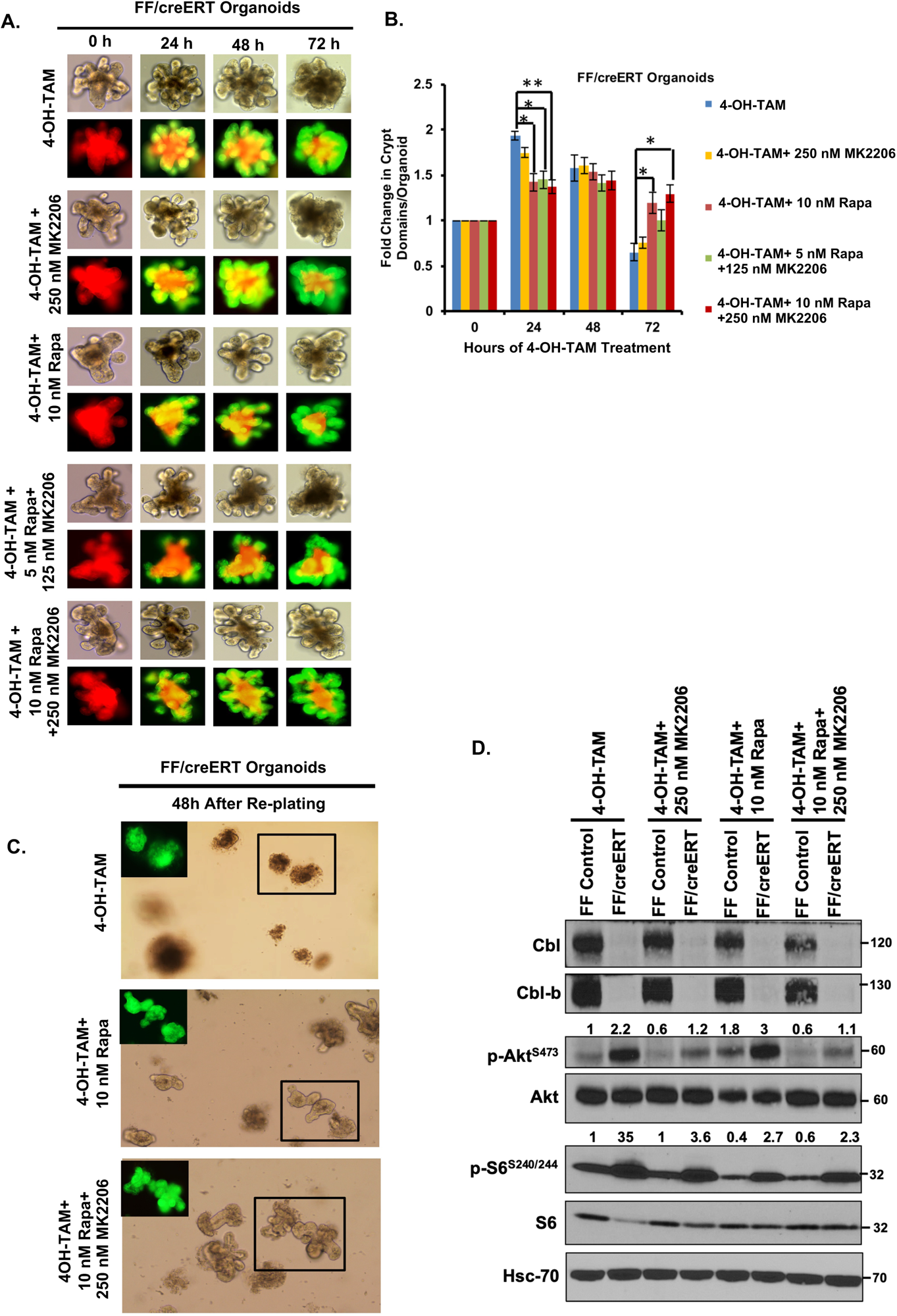
p-Akt/m-TOR pathway inhibition rescues defective maintenance of *Cbl/Cblb iDKO* intestinal crypt organoids. (A) Organoids established from FF/creERT mice carrying the mT/mG dual fluorescent reporter to document gene deletion were treated with 400 nM 4-OH-TAM, without (4-OH-TAM only control) or with Akt inhibitor MK-2206 (250 nM), mTOR inhibitor Rapamycin (10 nM) or their indicated combinations (5 nM Rapamycin plus 125 nM MK-2206 or 10 nM Rapamycin plus 250 nM MK-2206) and imaged at various time points over a 72 h period. While 4-OH-TAM alone treated organoid structures showed the expected disruption of architecture by 72 h, inhibition of Akt, mTOR or their combination rescued the organoid morphology with robust budding. Fluorescent imaging showed successful activation of Cre following TAM treatment under all conditions (conversion from uniform red to uniform green). (B) Quantification of crypt domains showed that Rapamycin by itself or in combination with MK-2206 significantly improved the crypt budding in comparison to TAM alone at 72 h; significant reduction in crypt budding was seen at 24 h, while no differences were seen at 48 h. (C) After 72h of organoid culture treatments, the 4-OH-TAM treated, 10nM Rapamycin+4-OH-TAM treated and 10 nM Rapamycin+250nM MK2206+4-OH-TAM treated organoids (the latter two showing the best rescue) were re-plated in complete ADF medium and imaged at 48 h after re-plating. The organoids previously treated with Rapamycin or Rapamycin plus MK-2206 showed robust budding while such structures were absent in secondary cultures of 4-OH-TAM alone treated organoids. (D) Western blot analysis of FF control organoids and FF/creERT organoids at 72 h of treatment showed the expected downregulation of p-Akt, p-S6 or p-S6 + p-Akt levels with 4-OH-TAM+MK-2206, 4-OH-TAM+ Rapamycin or 4-OH-TAM+MK-2206+Rapamycin treatment, respectively, as compared to 4-OH-TAM alone treatment in both FF Control and FF/creERT organoids. Densitometries of p-Akt/Akt and p-S6/S6 after normalizing to loading control in comparison to 4-OH-TAM treated FF control sample are indicated on the top of p-Akt and p-S6 bands. Quantified data are presented as mean +/- SEM of three independent experiments with statistics using *student’s two-tailed t-test*. ns, p ≤ 0.05, *; p ≤ 0.01, **.

## DISCUSSION

In addition to serving as a critical environment-organism interface with essential digestive and absorptive functions, the intestinal epithelium provides an elegant system to unravel gene functions critical for the maintenance and regeneration of epithelial tissues. Tyrosine kinase signaling linked to growth factor receptors represents one component of signaling mechanisms that orchestrate intestinal stem cell maintenance, proliferation, and differentiation (Clevers 2013; Tan and Barker 2015). Cbl-family ubiquitin ligases are activation-dependent negative regulators of tyrosine kinases but their roles in intestinal epithelial stem cell homeostasis are unknown. Here, using unique genetic deletion approaches, we establish a novel and redundant requirement for two Cbl-family members, Cbl and Cbl-b, in the maintenance of Lgr5+ ISCs and in their commitment to differentiate along various mature lineages. We also demonstrate the importance of negative regulation of the Akt-mTOR pathway by Cbl and Cbl-b as a critical mechanistic underpinning for their role in ISC regulation.

While *in vitro* studies of CBL family proteins have defined their key potential roles as negative regulators of both receptor and non-receptor tyrosine kinases, their *in vivo* physiologic roles have been primarily defined in the context of immune and hematopoietic cell compartments (Rao et al., 2002; Rathinam et al., 2008; Naramura et al., 2010; An et al., 2015; Goetz et al., 2016; Liyasova et al., 2015; Mohapatra et al., 2013: Tang et al., 2019; Li et al., 2019), with only sporadic studies of other physiological pathways and organ/tissue systems despite a pervasive role of tyrosine kinase signaling in physiological systems (Miettinen et al., 1995; Sibilia & Wagner, 1995; Batlle et al., 2002; Freier et al., 2005; Holmberg et al., 2006; Cheung et al., 2009; Vidrich et al., 2009; Sato et al., 2011; Al Alam et al., 2015). Notably, studies of the roles of Cbl proteins in epithelial biology are distinctly lacking except for our recent findings that Cbl and Cbl-b are vital for the post-natal development of mouse mammary gland, with their combined deletion leading to reduced ductal branching and defects in mammary stem cell self-renewal (Mohapatra et al., 2017). Given the counterintuitive nature of these findings in the face of a linkage of hyper-active tyrosine kinase signaling to breast and other epithelial cancers (Korkaya et al., 2008; Hynes and Watson, 2010; Radinsky et al., 1995; Guo et al., 2006; Martinelli et al., 2009), and a meager understanding of mammary epithelial stem cell maintenance and commitment into mature cell types or the signaling pathways involved (Sternlicht et al., 2006; Malhotra et al., 2011), the broader significance of Cbl proteins in epithelial stem cell maintenance has remained unclear. Thus, the current studies establishing a requirement for Cbl and Cbl-b in the maintenance of intestinal epithelial stem cells represent a significant step forward in our understanding of the biology of Cbl proteins, while expanding our understanding of the nature and importance of negative regulation of tyrosine kinase signaling in intestinal epithelial stem cell homeostasis.

Since no prior studies have systematically examined the expression or function of Cbl and Cbl-b proteins in the intestinal epithelium, we first assessed their expression pattern in the intestinal epithelium. We show that Cbl and Cbl-b are expressed throughout the length of the intestinal epithelial crypt-villus axis, but are enriched in the crypt (Fig. 1C, D), which pointed to their potential role in regulating the intestinal stem/progenitor cell compartment well-established to reside within crypts (Barker et al., 2007). Their overlapping expression pattern also suggested their potentially redundant functional roles in the intestine, reminiscent of their redundancy in regulating hematopoietic (Naramura et al., 2010; An et al., 2015) and mammary (Mohapatra et al., 2017) stem cells. Our findings of compensatory upregulation of Cbl expression in the crypts of *Cblb-*null mice and of Cbl-b in the villi of *Cbl-*null mice supported such redundant roles. In contrast to Cbl and Cbl-b, the expression of Cbl-c, measured indirectly using a *LacZ* reporter incorporated into the *Cblc* gene in *Cblc-*null mice, was confined to villi (Supplementary Fig. S1A). The lack of a compensatory increase in Cbl-b expression in the crypts of *Cbl-*null mice suggested a potentially more significant role for Cbl. Indeed, *Cbl-*null mice showed increased cell cycle entry within the crypt, goblet cell hyperplasia and reduction in the number of Paneth cells, while *Cblb-*null mice showed a mild increase in goblet cell numbers only (Fig.1C-D; Supplementary Fig. S3).

Since whole body *Cbl/Cblb* DKO mice are embryonic lethal (Naramura et al., 2002), we utilized the Lgr5+ ISC-specific inducible DKO of *Cbl* and *Cblb* (Mohapatra et al., 2017) to explore the redundant functional roles of Cbl and Cbl-b in ISCs. The *Cbl/Cblb* iDKO led to a rapid and marked drop in the size of the LgR5+ ISC compartment (Fig. 2D-G; Supplementary Fig. S2), with a concomitant increase in the Lgr5-Lo population, suggesting that Cbl and Cbl-b negatively regulate the commitment of ISCs into transit-amplifying progenitors, known to be Lgr5-Lo (Munoz et al., 2012; Basak et al., 2014). Supporting this conclusion, the iDKO mouse intestinal crypts exhibit increased cell proliferation, with no alterations in the rate of apoptosis (Fig. 3A-H); a similar but less pronounced crypt hyper-proliferation was seen in *Cbl^-/-^* mice (Fig. 1E, F). Furthermore, *Cbl/Cblb* iDKO was associated with crypt fission (Fig. 4G, H), a process that can be fueled by increased progenitor cell numbers (Dekaney et al., 2007; Barker 2014). Thus, our results support a role for Cbl and Cbl-b in the maintenance of a self-renewing pool of Lgr5+ ISCs by controlling the rate at which they commit to differentiation vs. self-renewal. Despite the hyper-proliferative crypt phenotype of *Cbl/Cblb* iDKO mice, however, a long-term hyperplasia of the epithelium or adenoma formation was never seen, a phenotype quite distinct from that seen in mice with deletion of the RTK negative regulator Lrig1, which induced epithelial hyperplasia and adenomas (Powell et al., 2012), even though Lrig1 is thought to function partly by facilitating RTK targeting by Cbl (Gur et al., 2004). Notably, although Lrig1 is expressed in Lgr5+ ISCs, its expression in the intestinal epithelium appears to be broader (Wong et al., 2012). The phenotype of Lrig1 deletion specifically in Lgr5+ cells has not been determined.

The lack of hyperplasia or adenomas upon *Cbl/Cblb* deletion in Lgr5+ ISCs, despite an immediate crypt hyper-proliferation phenotype, is likely to arise from multiple factors. Lineage-tracing showed a rapid transition of Lgr5+ cells with iDKO to enterocytes and goblet cells, and Goblet cell numbers were increased in *Cbl* or *Cblb* single KO mice and much more so in iDKO mice (Fig. 4A-D; Fig 1G-I). BrdU labeling showed proliferation zones to be limited to crypts (Fig. 3A), suggesting that the expanded transit-amplifying population must rapidly exit the cell cycle as the cells mature into differentiated cell types. We contend that the expansion of the transit-amplifying Lgr5-Low population occurs at the expense of the Lgr5-Hi compartment, which eventually eliminates all hyper-proliferating iDKO cells. Supporting that notion, lineage-tracing showed a decrease in the proportion of crypts with single blue (iDKO) cells (Fig. 4A-D), which are thought to represent a population of quiescent Lgr5+ cells that directly adopt a secretory cell fate, primarily into Paneth cells (Buczaki et al., 2013; van Es et al., 2012). Indeed, the numbers of Paneth cells were reduced in iDKO crypts (Supplementary Fig. S3A, B). A reduction in Paneth cell numbers could in turn further reduce the ISC numbers due to deficient niche function (Sato et al., 2011; Yilmaz et al., 2012; Geiser et al., 2012; Parry et al., 2013). The quiescent and cycling Lgr5+ pools are known to be in equilibrium (Takeda et al., 2011); apparently, the iDKO quiescent ISCs may be defective in resuming cycling Lgr5+ stem cell fate, and instead differentiate into progeny cells, thereby disrupting the quiescent-cycling ISC equilibrium. Interestingly, it has been previously shown that signaling downstream of RTKs can tilt the balance among different secretory cell types (Heuberger et al., 2014; Vidrich et al., 2009; Al Alam et al., 2015). A decrease in the quiescent Lgr5+ iDKO ISCs is consistent with the loss of quiescence in HSCs upon *Cbl/Cblb* deletion, but the shrinkage of the Lgr5-Hi cycling ISCs contrasts with an expansion of cycling HSCs (An et al., 2015). Whether this reflects a significant loss of Paneth cell niche in iDKO mice, rapid maturation of quiescent Lgr5+ cells along alternate fates (such as goblet cells or enterocytes) or a defective interconversion between the two different states of ISCs will require further studies.

It is noteworthy that the impact of *Cbl/Cblb* iDKO on mature intestinal epithelial cell types was not uniform, with a remarkable increase in goblet cells and reduction in Paneth cells providing a sharp contrast. Molecular marker analyses also suggested an increase in enterocytes with no change in enteroendocrine cells. These findings support the idea that Cbl/Cbl-b mediated regulation of ISC signaling is a determinant of differential fates of ISCs along various lineages. Consistent with altered fates of iDKO ISCs, the expression of markers of quiescent secretory progenitors, *Dll1* and *Ngn3*, was elevated in Lgr5-Hi cells while it was reduced in Lgr5-Lo cells of iDKO mice (Fig. 4E-F). High expression of *Dll1*, a ligand for Notch receptors, is a characteristic of +5 crypt cells, immediate descendants of Lgr5 ISCs that adopt a secretory cell fate (Stamataki et al., 2011; van Es et al, 2012). These cells are also thought to stochastically turn off Notch signaling and exit the cell cycle, further promoting their secretory cell fate. Notably, inhibition of Notch signaling is known to reduce the Lgr5-Hi cell population and the expression of stemness-associated genes such as *Lgr5* and *Olfm4* (Carulli et al., 2015; VanDussen et al, 2012) Ectopic expression of Notch in the Ngn3+ enteroendocrine progenitor cells, on the other hand, has been shown to drive differentiation towards enterocyte and goblet cell lineages (Li et al., 2012). Thus, it is possible that iDKO Lgr5-Lo cells experience greater than usual Notch signals propelling their expansion and further differentiation along enterocyte and goblet cell lineages. Further studies are needed to understand the basis of the differential impact of *Cbl/Cblb* iDKO on mature intestinal epithelial cell populations.

While *Cbl/Cblb* iDKO led to a rapid loss of Lgr5+ ISCs, no overt effects on mice were observed. One plausible reason for this is the replacement by ISCs without gene deletion since Lgr5-Cre expression is mosaic (Barker et al., 2007). To experimentally reveal a potential functional impact of the loss of Lgr5+ cells induced by *Cbl/Cblb* iDKO, we used a radiation injury model as recovery from such injury involves a rapid conversion of quiescent ISCs to cycling Lgr5+ ISCs (Takeda et al., 2011; Yan et al., 2012; van Es et al., 2012; Buczaki et al., 2013). Indeed, the *Cbl/Cblb* iDKO following focused beam abdominal radiation showed significantly slower recovery and re-establishment of the stem/progenitor compartments in the intestine (Fig. 5).

Since many growth factors and their cognate receptors have been linked to the maintenance of ISCs (Clevers 2013; Barker and Tan 2015) and signaling downstream of these could be impacted by *Cbl/Cblb* iDKO, we used unbiased single-cell RNA-seq analysis of control vs. iDKO intestinal organoids to discern the likely signaling pathways since Lgr5+ ISCs are known to divide, self-renew, and differentiate into mature cell types in organoid culture (Sato et al., 2009). Using a fully inducible DKO model (with tamoxifen-inducible CreERT and floxed *Cbl/Cblb*), our analyses revealed upregulation of the Akt/mTOR signaling pathway in iDKO ISCs and other cell compartments (Fig. 6J), findings that were confirmed by elevated pAKT and pS6 levels in iDKO organoids (Fig. 7K) and an increase in pS6 staining in ISCs of iDKO mice (Fig. 6L). The likelihood of the AKT-mTOR pathway being mechanistically important for ISC regulation by Cbl/Cbl-b is further strengthened by prior studies showing this axis to be a major target of Cbl/Cbl-b-imposed negative regulation in various systems (Li et al., 2009; Li et al., 2011; An et al., 2015) and our recent implication of the hyper-active AKT-mTOR pathway in MaSC depletion induced by loss of Cbl/Cbl-b (Mohapatra et al., 2017). Reversal of the loss of crypt domain-forming ability upon *Cbl/Cblb* iDKO by inhibiting Akt and/or mTOR in the organoid system (Fig. 7) further support the conclusion that negative regulation of Akt-mTOR signaling by Cbl/Cbl-b is required to maintain functional ISCs.

We posit that Cbl/Cbl-b provide a redundant but necessary negative regulatory mechanism to fine tune the level of Akt-mTOR signaling in Lgr5+ ISCs to favor self-renewal with orderly differentiation, while hyper-active Akt-mTOR signaling leads to rapid transition of ISCs into progeny at the cost of stem cells and aberrant lineage differentiation. mTOR is a major regulator of proliferation and growth during development as well as in adult tissues in mammals (Wullschleger et al., 2006), and plays its vital roles by integrating diverse signals including nutrient status, growth factor signals and redox/mitochondrial stress to control cellular metabolism. As such, mTOR signaling has emerged as critical in the control of self-renewal and differentiation in many stem cell systems, including the intestine (Ochocki and Simon, 2013). Overexpression of dominant negative TSC2, the negative regulator of mTOR, was shown to induce crypt proliferation but reduce goblet and Paneth cells by activating the Notch pathway (Zhou et al., 2015). On the other hand, inducible mTOR deletion in the intestine led to replacement of mTOR-deficient ISCs by WT ISCs while constitutive mTOR deletion led to altered differentiation, including blunted villi, loss of alkaline phosphatase, and loss of Goblet and Paneth cells (Sampson et al., 2016). In addition, mTOR-deleted crypts failed to form organoids *in vitro*. Interestingly, calorie restriction was shown to inhibit the mTOR pathway in Paneth cells and enhance their niche function to support ISC self-renewal, but exposure of both Paneth and Lgr5+ stem cells to Rapamycin enhanced organoid formation ability of the latter *in vitro* (Yilmaz et al., 2012). PTEN, a negative regulator of PI3-kinase-Akt signaling has been linked to the maintenance of a quiescent ISCs (He et al., 2007) and progenitors (Richmond et al., 2015). Loss of YY1, an effector of mTOR, has been shown to cause exhaustion of Lgr5+ stem cells (Perekatt et al., 2014). It is likely that the various reported results represent the distinct outcomes of mTOR signaling in different intestinal epithelial cell and ISC compartments. We also note that our current understanding of Cbl/Cbl-b as tyrosine-kinase directed ubiquitin ligases suggests that their negative regulatory role in Akt-mTOR signaling in ISCs reflects their activation downstream of tyrosine kinase-coupled receptors (Mohapatra et al., 2013). Whether Cbl/Cbl-b might also regulate the Akt-mTOR axis in response to nutritional cues remains to be determined. Future studies of perturbations in Akt-mTOR signaling pathways specifically in Lgr5+ ISCs with or without concurrent *Cbl/Cblb* deletion should further elucidate the importance of Cbl/Cbl-b regulation of the Akt-mTOR pathway in ISC maintenance.

Pertinent to our findings, previous studies reported that Cbl can serve as a negative regulator of beta-catenin through direct binding and ubiquitination (Chitalia et al., 2013; Shivanna et al., 2015). Subsequently, the same investigators reported an inverse correlation between Cbl expression and nuclear beta-catenin levels in colorectal tumors, with increased proliferation and tumorigenic ability of colorectal tumor cell lines with downregulation of CBL expression (Shashar et al., 2016; Kumardevan et al., 2018). Furthermore, haploinsufficiency of *Cbl* in mice enhanced the atypical hyperplasia and adenocarcinoma numbers in APC^Δ14/+^ mice but had no effect on its own (Richards et al., 2020). While we have not directly examined the WNT signaling pathway, we consider it unlikely that the aberrations of ISCs and mature intestinal epithelial cell lineages in *Cbl/Cblb* iDKO mouse model reflect a potential increase in beta-catenin activity since Lgr5+ ISC-specific increased WNT/beta-catenin pathway activation upon *Apc* deletion has been demonstrated to be tumorigenic (Barker et al., 2009).

Finally, it is notable that aberrant tyrosine kinase and Akt-mTOR signaling are linked to oncogenesis, including in colorectal cancer (Zenonos and Kypriano, 2013; Chen et al., 2015), yet somatic mutations of Cbl/Cbl-b found in myeloid leukemias (Caligiuri et al., 2007; Muramatsu et al., 2010; Kao et al., 2011; Jawhar et al., 2015) are distinctly rare in colorectal and other epithelial malignancies. Future studies to determine if the requirement for Cbl/Cbl-b for maintenance of stem cell compartments in normal epithelial tissues, such as the intestine (here) and mammary glands (Mohapatra et al., 2017), extends to cancer stem cells that drive these malignancies (Barker et al., 2009; Merlos-Suarez et al., 2011; Schepers et al., 2012; Pece et al., 2010) will therefore be of great interest.

## MATERIALS AND METHODS

### Mouse Strains

The Lgr5-EGFP-IRES-creERT2 knock-in (*Lgr5^tm1(cre/ERT2)Cle^*; Stock No: 008875) and Rosa26-LacZ reporter (B6.129S4-*Gt(ROSA)26Sor^tm1Sor^*/J; Stock No: 003474; aka R26R) mice were purchased from The Jackson Laboratory and interbred; this genotype served as a control. These mice were further bred with *Cbl^flox/flox^*; *Cblb^-/-^* mice (Naramura et al., 2010) to obtain *Cbl^flox/flox^*; *Cblb^-/-l^*; *Lgr5-cre^ERT2^*; *R26-LacZ* (iDKO) experimental mice for tamoxifen-inducible intestinal stem cell specific deletion of *Cbl* and *Cbl-b*. The *Cbl^-/-^* (Hua Gu et al.1998) and *Cblb^-/-^* (Chiang et al., 2000) mice used to examine the function of CBL and CBL-B individually have been reported previously. Age and gender matched wild-type (WT) C57BL/6J mice served as controls. *Cbl^flox/flox^; Cblb^flox/flox^* (Conditional *Cbl*/*Cblb* double floxed (FF control) mice generated in our lab (Goetz et al., 2016) were crossed to Rosa26-CreERT2 (B6.129-*Gt(ROSA)26Sor^tm1(cre/ERT2)Tyj^*/J; Stock No: 008463; aka R26-CreERT2; The Jackson Laboratory) and mT/mG-Reporter mice (B6.129(Cg)-*Gt(ROSA)26Sor^tm4(ACTB-tdTomato,-EGFP)Luo^*/J; Stock No: 007676; aka mT/mG, mTmG; The Jackson Laboratory) to obtain the *Cbl^flox/flox^*; *Cblb^flox/flox^*; *R26Cre^ERT2^*; *mT-mG* model used to isolate intestinal crypts for organoid culture studies. All mouse strains were maintained on a C57BL/6J background under specific pathogen-free conditions and handled in accordance with protocols approved by the Institutional Animal Care and Use Committee (IACUC) of UNMC. All mice used in this study were euthanized by first anesthetizing using isoflurane (Piramal Healthcare, Catalog no. NDC66794-017-25) followed by cervical dislocation to confirm death. Mice were genotyped using tail DNA PCR (primer sequences in Table-S1).

### Tamoxifen-induced Cre Activation

For *in vivo* deletion of floxed Cbl alleles, Tamoxifen free base (MP Biomedicals, Catalog no. ICN15673880) was resuspended in sunflower oil (Sigma Aldrich, Catalog no. S5007) at a concentration of 10 mg/ml. At 6 weeks of age, gender matched Lgr5-EGFP-IRES-creERT2; *R26-LacZ* (WT-Control) and *Cbl^flox/flox^*; *Cblb^-/-l^*; *Lgr5-cre^ERT2^*; *R26-LacZ* (iDKO) mice were given an intra-peritoneal injection of 1mg tamoxifen for 3 consecutive days to activate inducible Cre recombinase. Animals were analyzed 3 days after the last injection (5 days after the first injection), 8 days after the last injection (10 days after the first) or 4 months after the last injection. For *in vitro* activation of Cre, crypts were isolated from FF control and FFcreERT mice and equal numbers were plated in Matrigel (Corning, catalog# 356231) under conditions as described (Sato et al., 2011) to obtain organoids by day 7. The organoids thus generated were re-plated at 1:4 split ratio and exposed to 400 nM 4-hydroxy tamoxifen (Sigma; Catalog #H7904) in medium to achieve the deletion of *Cbl* and *Cblb*. For rescue experiments, FF control and FFcreERT organoids were treated with Rapamycin (Sigma, Catalog#R0395) and Akt-inhibitor, MK2206 2HCl (Selleckchem.com, Catalog#s1078) alongside 4-OH-tamoxifen.

### Tissue Harvest and Histological Analysis

Mouse small intestine was carefully excised from the peritoneal cavity by first snipping at the ileo-cecal junction and pulling away to disentangle the intestine from the mesh of mesentery and finally cutting at the pyloric end of stomach. The intestine was immediately placed in a Petri dish, and gently flushed with cold phosphate buffered saline (PBS) to remove fecal matter using a 30 ml syringe and a blunt ended needle. Once cleaned, intestine was placed on a blotting sheet, cut open longitudinally, swiss-rolled and placed in formalin (Sigma Aldrich, Catalog No. HT501128) for overnight fixation at room temperature. Fixed tissue was embedded in paraffin and sectioned to 5 μM thickness.

### Immunofluorescence (IF)

Tissue sections were deparaffinized and rehydrated followed by antigen retrieval in sodium citrate antigen unmasking solution (Vector Laboratories; catalog #H-3300-250) by heating in a microwave oven for 20 min. Slides were then washed in PBS and blocked for 1h in 20 mm Tris Buffered Saline with 0.1% Tween-20 (BioRad, catalog #1610781) (TBS-T) containing 5% goat serum (Sigma Aldrich, G9023) followed by overnight incubation at 4°C with optimized concentrations of primary antibodies [(BrdU, Developmental Studies Hybridoma Bank, Iowa, G3G4,1:25), Chromogranin A (Abcam, Catalog No.15160, 1:200), Cleaved Caspase 3 (CST, Cat# 9664, 1:200), phospho-histone 3 (Ser10) (Cat# 9701, CST, 1:200), GFP (CST, Cat# 2956, 1:200), GFP (Aves Labs, Chicken, Cat#GFP-1010, 1:2000), phospho-S6 (CST, Cat# 4858,dilution 1:200). Tissue sections were then washed thrice in PBS for 5 minutes each, followed by incubation with fluorescence-tagged secondary antibodies (Molecular Probes Alexa series, all 1:400 in PBS) for 1h at room temperature, washed 3X with PBS and mounted with Vectashield and DAPI to stain nuclei (Vector Laboratories).

### Immunohistochemistry (IHC)

Tissue sections were deparaffinized and rehydrated followed by antigen retrieval in sodium citrate antigen unmasking solution (Vector Laboratories) using microwaving for 20 min. Slides were then washed in PBS and incubated in 5% H2O2 for 30 minutes at room temperature to inactivate endogenous peroxidase. The slides were washed with PBS and incubated with the following blocking solutions for 1 hour at room temperature: Mouse Ig Blocking Reagent in Mouse on Mouse (MOM) kit from Vector Laboratories (PK-2200) for staining with antibodies against Cbl (mouse monoclonal, anti-Cbl, BD Bioscience, 1:400) and Cbl-b (mouse monoclonal, anti-Cbl-b, Abcam, 1:100); 5% goat serum (Sigma Aldrich, G9023) for staining with antibodies against Chromogranin A (1:200, Abcam) and Ki67 (Rat eBioscience, 145698-82, 1:200). Slides were washed twice with PBS and incubated with primary antibodies indicated above at 4°C overnight. The next day, slides were washed thrice with PBS for 5 min each and incubated with species-matched biotinylated secondary antibodies for 30 min. The ABC amplification IHC kit (Vector Laboratories, PK-6100) was used as per vendor’s instructions to amplify the signal. Slides were washed twice, and peroxidase-based signals were developed using DAB (3,3’-diaminobenzidine, Vector Laboratories, SK-4100) as substrate. The slides were counterstained with hematoxylin, dehydrated and mounted in automated counter-stainer (Sakura Tissue-tek Prisma) at the UNMC Tissue Science Core Facility. In all the cases, a species matched isotype IgG, genetic knockouts or secondary antibody only negative controls were used.

### **β**-Galactosidase (LacZ) Staining

The β-galactosidase (LacZ) staining for lineage tracing was performed in intestinal sections of mice with a lacZ reporter, as described (Barker and Clevers, 2010). Briefly, freshly dissected intestinal tissue was flushed clean with cold PBS and fixed in glutaraldehyde fixative (1% formaldehyde, 0.2% glutaraldehyde, 0.02% NP-40 in PBS) for 2 hours at 4°C. Tissue was washed twice with PBS and incubated in equilibration buffer (2 mM MgCl2, 0.02% NP-40, 0.01% sodium deoxycholate in PBS) for 30 minutes at room temperature. Subsequently, the tissue was incubated in LacZ substrate [5mM K3Fe(CN)6, 5mM K4Fe(CN)6, 2mM MgCl2, 0.02% NP-40, 0.1% sodium deoxycholate, 1 mg/ml X-gal in PBS] for 4 hours at 37°C, protected from light. Once the blue color developed, tissue was washed twice with PBS, swiss-rolled, and incubated in 4% paraformaldehyde at 4°C overnight. The fixative was then removed, and tissue was washed with PBS and dehydrated in alcohol gradient (70%, 96% and 100% for 1 hour each) and paraffin embedded. 5 μM tissue sections were cut, paraffin cleared and rehydrated, followed by either counterstaining with nuclear fast red (Vector Labs), PAS staining or IHC as indicated above.

### Intestinal epithelial stem cell (ISC) isolation and flow cytometric analysis

A modification of the Intestinal Stem Cell Consortium protocol (Wang et al., 2013) was used to obtain stem cell-enriched single cell suspensions of intestinal epithelium. Briefly, freshly dissected intestinal tissue was flushed clean with ice cold wash buffer (100 µg/ml penicillin/streptomycin, 10mM HEPES, 2mM Glutamax in HBSS) and cut open longitudinally to expose the lumen. A glass coverslip was used to scrape off the villi and the tissue chopped into 5 mm long pieces. The chopped pieces were then washed several times with cold wash buffer by vigorously pipetting until the supernatant was almost clear. After this, the tissue was placed in dissociation reagent #1 [15 mM EDTA, 1.5 mM DTT, 10 µM ROCK inhibitor (Y27632, TOCRIS), 100 µg/ml penicillin/streptomycin, 10mM HEPES, 2mM Glutamax in HBSS] and rocked slowly at 4°C for 30 minutes. Supernatant was removed and replaced with fresh cold wash buffer and the tubes were shaken vigorously. The tissue pieces were allowed to settle down, and the supernatant was passed through 70 µM cell strainers and centrifuged at 1200 rpm for 5 minutes at 4°C. The pellet was resuspended in 10 ml wash buffer and centrifuged at 600 rpm for 2 minutes at 4°C. The cell pellet obtained was resuspended in dissociation reagent # 2 [TrypLE Express (Thermo Scientific), 10µM ROCK inhibitor (Y27632, TOCRIS), 0.5 mM N-acetylcysteine (Sigma, Catalog#A7250), and 200 µg/ml DNase-I (Stem Cell Technologies, Catalog#07900) in Intestinal Epithelial Stem Cell Media (ISC Media) [Advanced DMEM/F12 supplemented with 1XN2 (Gibco, Catalog#17502048), 1XB27 without vitamin A (Gibco, Catalog#12587010), 10 mM HEPES, 2mM Glutamax (Gibco, Catalog#35050061),100 µg/ml penicillin/streptomycin] and incubated for 8 minutes at 37°C, with intermittent pipetting to reduced cell clumping. Equal volume of neutralization buffer (10% FBS in ISC Media) was added, and the cell suspension was passed through 20 µM cell strainers. The cells were then centrifuged at 1000 rpm for 5 minutes at 4°C. The supernatant was discarded, and cells were resuspended in 1 ml of cold ISC medium and cell numbers counted to assess the yield. The isolated cells were stained with Alexa 700 conjugated anti-CD45 antibody (Catalog No. 103128, 0.25 µg per 10^6^ cells) at 4°C for 30 min, washed twice and propidium iodide (5 µg per 10^6^ cells) added before FACS analysis on a BD LSR2 instrument. FACS data were analyzed using BD FACS DIVA software.

### RNA Isolation, cDNA synthesis and quantitative PCR analysis

Cell analyzed by flow cytometry were directly sorted into 500 µl of RNA lysis buffer of the RNAqueous®-Micro Total RNA Isolation Kit from Thermo Scientific (Catalog No. AM1931). Instructions from the kit were followed to isolate the RNA. The RNA yield was assessed using a Nanodrop instrument and cDNA was synthesized using a kit from QIAGEN (catalog No. 205311). QuantiTect SYBR® Green PCR Kit was used to perform real-time PCR analysis on a Bio-Rad CFX-96 Thermo Cycler. Sequences of primers (obtained from Sigma-Aldrich) are listed in Table-S2. Relative gene expression was calculated according to 2ΔΔCt method.

### Droplet-based scRNA-seq

Single cell gene expression profiling was performed as per manufacturer’s protocol (10X genomics, Pleasanton, CA) by the UNMC Genomics core facility. Briefly, intestinal crypts were isolated from two 10-week-old *Cbl^flox/flox^*; *Cbl-b^flox/flox^*; *R26Cre^ERT2^* mice and grown into intestinal organoids using our established protocol. The organoids were replated and grown in presence or absence of 4-hydroxy-tamoxifen (400 nM) for 3 days to delete *Cbl* and *Cblb.* Organoids were dissociated into single cells. The single cells were checked for viability and processed through the GemCode Single Cell Platform. An input of 7,000 cells was added to each channel of a chip with a recovery rate of 1,200 cells on average. The cells were then partitioned into Gel Beads in Emulsion (GEMs) in the GemCode instrument, where cell lysis and barcoded reverse transcription of RNA occurred, followed by amplification, shearing and 5′ adaptor and sample index attachment. Libraries were sequenced on an Illumina NextSeq 500 in the Genomic Core facility, UNMC. Demultiplexing, alignment to the transcriptome and UMI-collapsing were performed through 10X genomics recommended cell ranger standard pipeline (Cell Ranger toolkit, version 3.0.1) by the Bioinformatics and Systems Biology Core at UNMC. Seurat was used to discard low quality cells with aberrant gene expression, normalization, identification of highly variable features, scaling and linear dimensionality reduction. Subsequently clustering the cells was carried out by t-distributed stochastic neighbor embedding (t-SNE) method (Butler et al., 2018). Cluster markers were then identified in Seurat by inspecting differentially expressed features among the clusters (Haber et al., 2017). Signaling pathways were analyzed using Gene Set Enrichment Analysis (GSEA). The gene set enrichment score was calculated using the Bioconductor GSVA package and the plots were generated using R.

### Lysate preparation and Western Blot

Freshly-excised, cleaned and chopped fragments of intestine were incubated in 2 mM EDTA in cold PBS at 4°C for 30 minutes with vigorous shaking to detach the intestinal epithelium from the remaining tissue. Once the epithelial layer was separated, the supernatant containing the detached epithelium was pelleted and homogenized in RIPA lysis buffer containing protease and phosphatase inhibitors. Intestinal organoid lysates were also prepared in the same lysis buffer. The lysate proteins were quantified using the BCA kit from Thermo Scientific. 40 µg of lysate protein aliquots were resolved on 8% SDS–PAGE and transferred to PVDF membranes, which were then blocked with 5% BSA in TBS-T and incubated overnight at 4°C with specific antibodies diluted in TBS-T. The following antibodies were procured from commercial sources: Anti-Cbl (Clone 17/c-Cbl, BD Bioscience); anti-Cbl-b (Clone D3C12, Cell Signaling Technology); anti-HSC70 (Clone B6, Santa Cruz Biotechnology Inc. Santa Cruz, CA), phospho Akt (Ser473) (Cat# 4060, Cell Signaling Technology); Akt (pan, Cat# 4691, Cell Signaling Technology); phospho S6 (Ser240/244), Cat# 5364, Cell Signaling Technology); Total S6 (5G10, Cat#2217, Cell Signaling Technology), phospho-histone 3 (Ser10) (Cat# 9701, Cell Signaling Technology), and Villin-1 (Cat# 2369, Cell Signaling Technology). Membranes were washed 3X in TBS-T, incubated with HRP-conjugated species-specific secondary antibodies (Zymed Laboratories) and signals were detected with Pierce ECL substrate (Thermo scientific).

### Focused Beam Abdominal Radiation

Mouse irradiation was accomplished on the TrueBeam^TM^ STx linear accelerator (Varian Medical Systems, Palo Alto, CA) in the Radiation Oncology Department at UNMC. Mice were first sedated with Ketamine/Xylazine, restrained to prevent movement to ensure reproducible positioning for radiation and simulated using the simulation CT scanner. The radiation target abdominal region was contoured by an attending radiation oncologist. A specific treatment plan was created for each mouse by the medical physicists, and mice were treated to a dose of 14 Gy photon radiation.

### Organoid Culture

A modification of the organoid protocol established by Sato et al. (Sato et al., 2011) was used to prepare organoid cultures from crypts of *Cbl^flox/flox^*; *Cblb^flox/flox^*; *R26Cre^ERT2^* mice. Freshly dissected, cleaned and chopped fragments of ileum were incubated in 2 mM EDTA in cold PBS at 4°C for 30 minutes in a 15-ml Falcon tube. Tissue fragments were allowed to settle down, the supernatant was replaced with wash buffer (described under isolation of ISC) and the tube was shaken by hand for 3 minutes to release the epithelial layer from the remaining tissue. The supernatant was passed through 70 μm cell strainer and the strainers further rinsed with fresh wash buffer. The crypts were counted under the microscope and aliquots containing 500 crypts was pelleted. The supernatant was removed, and the crypt pellets were resuspended in Matrigel (BD Bioscience, Catalog No. 356237) mixed with growth factors (EGF 50 ng/ml; Noggin 100ng/ml; R-spondin 500 ng/ml) and plated in wells of 24 well plates (Costar). The plates were kept at 37°C for 15 minutes to allow the Matrigel to polymerize. 500 μl of ISC medium was added and plates were incubated at 37°C in 5% CO2 tissue culture incubators. On the 4^th^ day after plating, the spent medium was replaced with ISC medium containing EGF, Noggin, R-spondin1 conditioned medium (1:10) and Wnt-3a conditioned medium (1:10). The conditioned media were obtained from late log phase cultures of HEK293T cells overexpressing recombinant R-spondin1 (a gift from Dr. Mark R. Frey, Keck School of Medicine, University of Southern California) or L cells overexpressing Wnt-3a (ATCC).

### Fractionation of mouse crypt and villus

Biochemical fractionation of mouse crypt and villus was carried out as described by Mahmoudi et al. (Mahmoudi et al., 2009). Briefly, freshly dissected small intestine of 12-week-old wildtype, *Cbl^-/-^* and *Cblb^-/-^* mice were cleaned and flushed with ice-cold PBS several times and cut open longitudinally to expose crypts and villi. Small intestine was chopped into small 1–2 cm fragments and incubated in 2mM EDTA in cold Ca^2+^/Mg^2+^ free PBS (Chelation buffer) at 4°C for 20 minutes. Tissue fragments were allowed to be settled down and the supernatant was replaced with fresh chelation buffer and the tube was shaken vigorously for 10 minutes to release the epithelial layer from the remaining tissue. This process was repeated, and the fractions were passed through 70 μm strainer. The villus structures remain on top of the cell strainer and cells were collected, whereas the flow through was discarded. The process of incubation, shaking and separation through the cell strainer was repeated. The fractions 2-4 was collected for pure villi. The intact crypts remain in flow through after fifth fraction. The purified villi and crypts were washed with cold PBS and protein lysates were prepared.

### Statistical Analyses

Quantified results of LacZ stained sections, qPCR, immunohistochemistry, and flow cytometry were compared between groups using the Student’s t-test on GraphPad Prism, and were presented as mean +/- SEM, with p ≤0.05 deemed as significant.

## ACKNOWLEDGEMENTS

We thank the Band Lab members for their many discussions, Drs. Rakesh Singh and Karen Gould for advice, and the staff of UNMC Core facilities for their invaluable assistance. We thank Dr. Mark R. Frey, Keck School of Medicine, University of Southern California for providing the Rspondin-1 secreting cell line. This work was supported by the National Institutes of Health (CA105489, CA99163 to H.B); the Nebraska Department of Health and Human Services LB606 (18123-Y3 to H.B.); a pilot grant from the UNMC Regenerative Medicine Program; a Core Facility usage grant for RNA-seq analysis from the Nebraska Research Initiative; and by support to the UNMC Confocal, Flow Cytometry, Bioinformatics, Molecular Biology, and other Core facilities from the NCI Cancer Center Support Grant to the Fred & Pamela Buffett Cancer Center (P30CA036727) and the Nebraska Research Initiative. We thank the Dr. Raphael Bonita Fund for partial support of graduate students in the H.B. lab. N.Z., B.C.M., and W.A. received University of Nebraska Medical Center graduate assistantships. B.T.G. was a trainee under the National Cancer Institute Cancer Biology Training Grant T32CA009476.

