## Supplemental data for "Cbl and Cbl-b Ubiquitin Ligases are Essential for Intestinal Epithelial Stem Cell Maintenance"

### Supplementary Information

#### Supplementary Tables and Figures

| Supplementary Table S1. List of primers used for PCR-based genotyping of various alleles | | |
|---|---|---|
| Gene | Forward 5'-3' | Reverse 5'-3' |
| *Cbl* WT | AAGTTCCAAGCCTAGCCAGATA TGTGTGTG | TCCCCTCCCCTTCCCATGTTTTTAATAGACTC |
| *Cbl* Del | TGGCTGGACGTAAACTCCTCTT CAGACCAATAAC | TCCCCTCCCCTTCCCATGTTTTTAATAGACTC |
| *Cbl* floxed | GTGGTGGCTTGCAATTATAATC CTACCACTTAGG | GTTTGAGATGTCTGGCTGTGTACACGCG |
| *Cblb* del | CCCAGCAAAAGTAGCCAATG | CTTGCAAAAAGGACTAAGATTC |
| *Cblb* floxed | GGCAGAACCACTGAGACACATTTA | GGCTGCCAAACTGCTACCCAGGAG |
| Lgr5 - CRE ERT2(WT) | CTGCTCTCTGCTCCCAGTCT | ATACCCCATCCCTTTTGAGC |
| Lgr5- CRE ERT2 (Mutant) | CTGCTCTCTGCTCCCAGTCT | GAACTTCAGGGTCAGCTTGC |
| Cre | GCGGTCTGGCAGTAAAAACTATC | GTGAAACAGCATTGCTGTCACTT |
| R26-LacZ (WT) | AAA GTC GCT CTG AGT TGT TAT | GGA GCG GGA GAA ATG GAT ATG |
| R26-LacZ (Mutant) | AAA GTC GCT CTG AGT TGT TAT | GCG AAG AGT TTG TCC TCA ACC |

| **Supplementary Table S2. List of real time qPCR primers used to detect mRNA levels of the indicated genes** | | |
|---|---|---|
| Gene | Forward 5’-3’ | Reverse 5’-3’ |
| *Cbl* | AGCTGATGCTGCCGAATTT | TTGCAGGTCAGATCAATAGTGG |
| *Cblb* | GGAGCTTTTTGCACGGACTA | TGCATCCTGAATAGCATCAA |
| *Gapdh* | CCTGGAGAAACCTGCCAAGTATG | AGAGTGGGAGTTGCTGTTGAAGT |
| *Dll1* | CCCATCCGATTCCCCTTCG | GGTTTTCTGTTGCGAGGTCATC |
| *Nurog3* | GAGTCGGGAGAACTAGGATG | CAGTCCCTAGGTATGAGAGT |
| *Lgr5* | AACGGTCCTGTGAGTCAACC | ATGGGGTAAGCTGGTGATGC |
| *Olfm4* | ACTTTCCAATTTCACTGGGT | GGACATACTCCTTCACCTTAG |

**Supplementary Fig. S1**

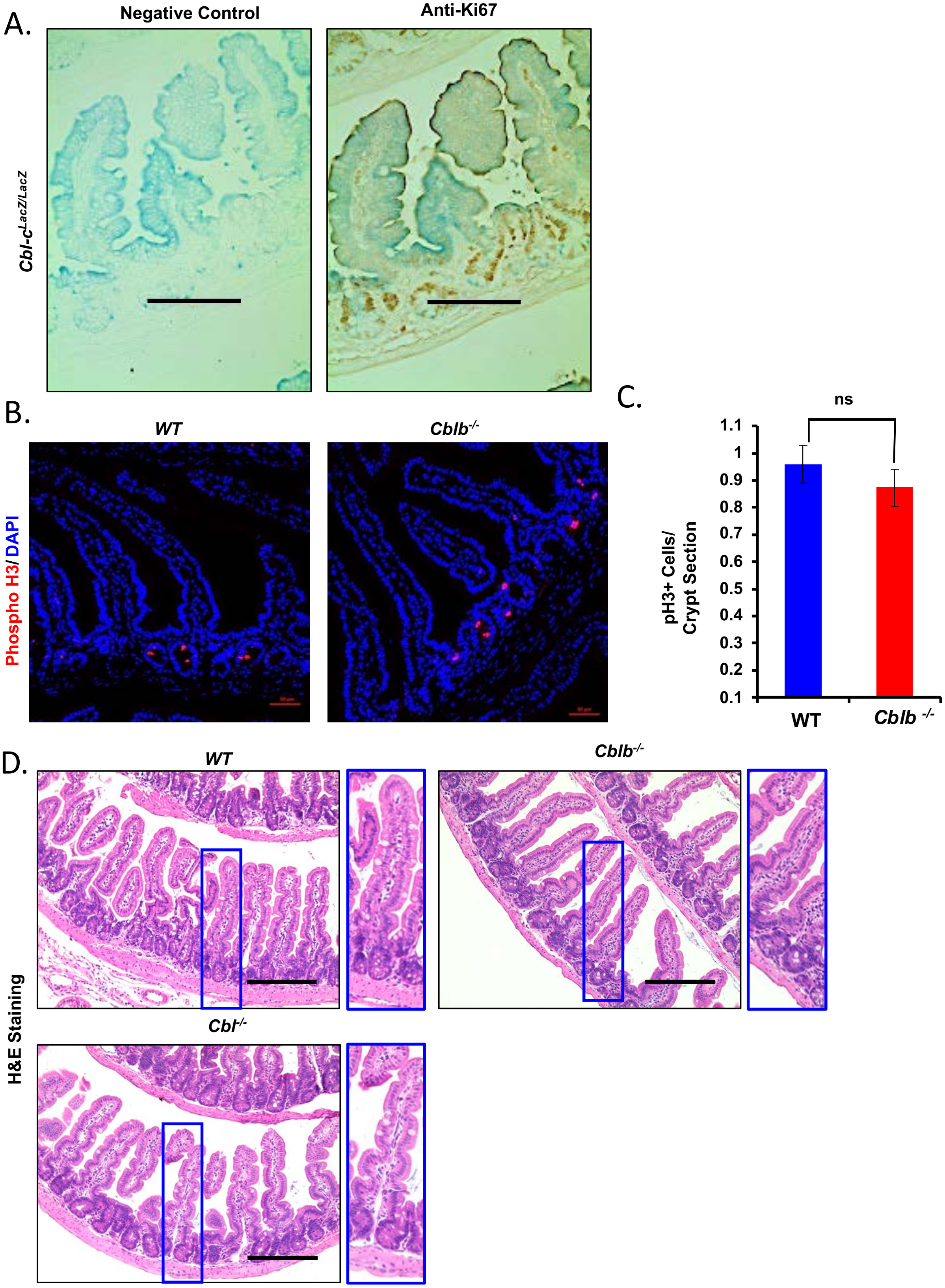

**Supplementary Fig. S1. Expression of Cbl proteins in the intestine and impact on epithelial architecture.** (A) Cbl-c expression was examined using mice with a *LacZ* expression cassette knocked-in in the *Cblc* gene and expressed from the *Cblc* promoter (*Cblc*$^{LacZ/LacZ}$, Griffiths et al., 2003). FFPE sections of the intestine from these mice were co-stained for X-gal (blue) and Ki67 (brown). Staining with the secondary anti-rat antibody alone was used as a negative control. Intense blue staining, reflecting *Cblc* expression pattern, was predominantly seen within the villi, while brown Ki67 staining marked the crypt compartment with little or no blue staining. (B) Immunofluorescence staining using anti-phospho-histone 3 (pH3) antibody (red) was performed on age-matched *WT* vs *Cblb*$^{-/-}$ mice to evaluate any changes in the proportion of proliferative cells; DAPI (blue) was used to stain the nuclei, scale bar = 50 µm. (C) Quantitative analysis of results presented in B showed no significant difference in the number of pH3+ cells per crypt between the two genotypes. (D) Hematoxylin & Eosin staining of WT, *Cbl*$^{-/-}$, and *Cblb*$^{-/-}$ mouse intestinal sections reveled no structural defects in the intestines of single gene knockouts. Scale bar=200 µm. Quantified data are presented as mean +/- SEM of 3 experiments using *student's two-tailed t-test***.** ns, $p \le 0.05$, *; $p \le 0.01$, **; $p \le 0.001$, ***.

**Supplementary Fig. S2**

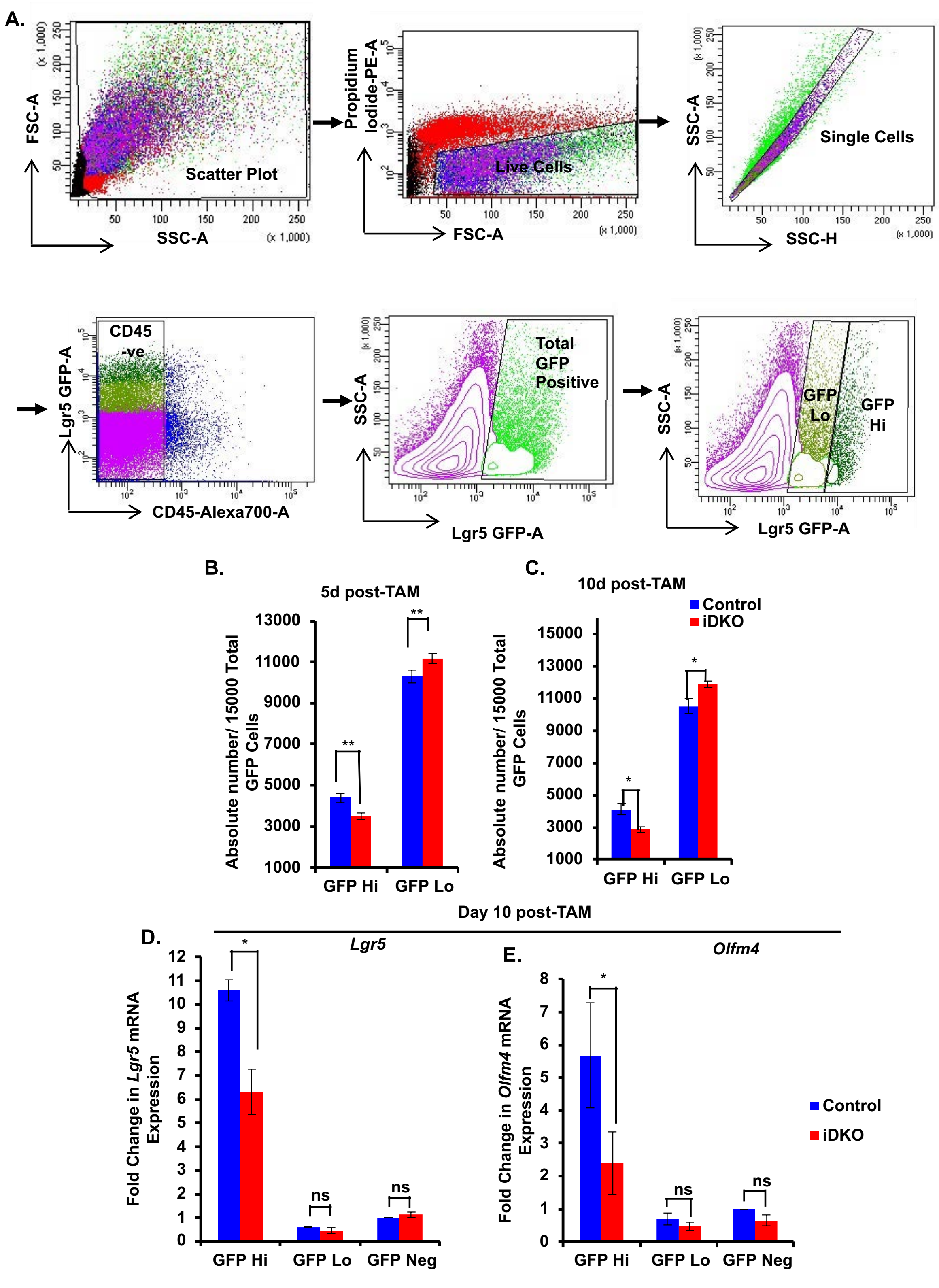

**Supplementary Fig. S2. Expansion of intestinal progenitors with loss of the stem compartment upon Lgr5+ cell-specific *Cbl/Cblb* iDKO.** (A) FACS analysis was performed to assess the impact of *Cbl/Cblb* deletion on *Lgr5*+ ISCs and their immediate progeny. The gating strategy shown in A employed scatter gate, propidium iodide, doublet discrimination gate and CD45 staining to exclude debris, dead cells, doublets and non-epithelial cells, respectively. *Lgr5-cre* negative samples were included each time to set the gate for total GFP-positive population. A contour plot of the total GFP pool was used to assign GFP Hi (stem cells) and GFP Lo (progenitors) subpopulations. (B, C) Quantification of the absolute numbers of GFP expression-based subpopulations showed a significant drop in stem cells and a concomitant increase in progenitor cells in *Cbl/Cblb* iDKO mice as compared to controls at both day 5 (B) and day 10 (C) post-TAM induction. (D, E) Quantitative real-time PCR was performed for stemness associated genes, *Lgr5* (D) and *Olfm4* (E) on GFP Hi, GFP Lo and GFP negative sorted fractions from control and *Cbl/Cblb* iDKO mice. Both genes showed the expected enrichment in the GFP Hi pool in both genotypes, validating the gating strategy. A significant reduction in the expression of *Lgr5* and *Olfm4* genes is seen in the GFP Hi compartment of *Cbl/Cblb* iDKO mice. Quantified data are presented as mean +/- SEM of 4 independent experiments with statistics using *student's two-tailed t-test***.** ns, $p \leq 0.05$, *; $p \leq 0.01$, **.

### Supplementary Fig. S3

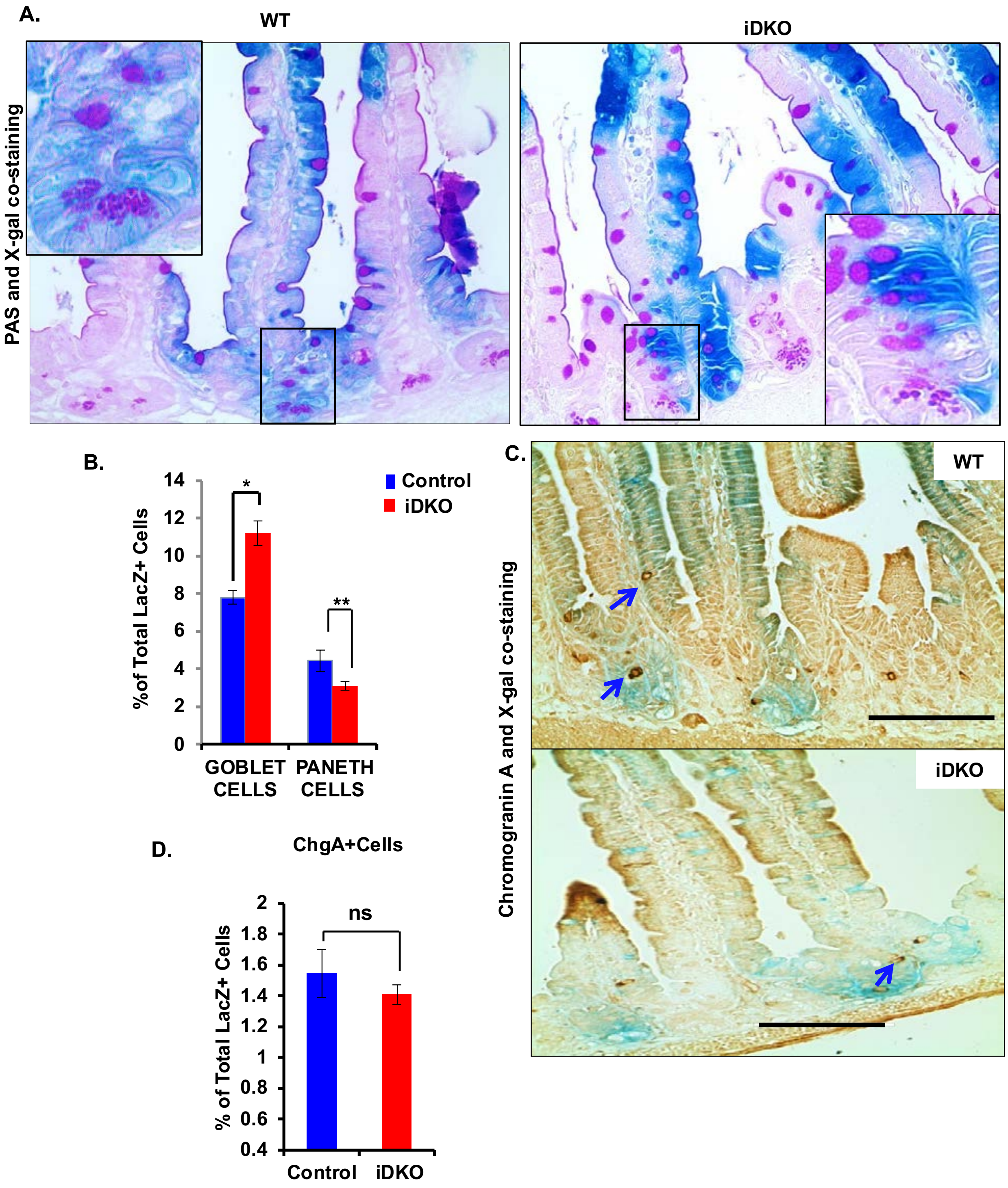

**Supplementary Fig. S3. Altered intestinal epithelial cell differentiation upon Lgr5+ cell-specific *Cbl/Cblb* iDKO in the intestine.** Control and *Cbl/Cblb* iDKO mouse intestinal sections were serially stained with X-gal followed by PAS to visualize goblet and Paneth cells (A), or with X-gal followed by Chromogranin A (ChgA) (C) to visualize enteroendocrine cells. (A) Representative images of X-gal plus PAS staining. (B) Two thousand *LacZ*+ cells were counted from jejunum and ileum regions per animal and the number of PAS+ goblet and Paneth cells that co-stained with X-gal were enumerated. Quantification shows a significant increase in the % of *LacZ*+ PAS+ goblet cells and a significant decrease in the % of *LacZ*+ PAS+ Paneth cells in *Cbl/Cblb* iDKO animals as compared to controls. (C) Representative images of X-gal plus ChgA staining. (D) Quantification of ChgA-LacZ dual positive cells revealed no significant difference in the % of ChgA-LacZ dual positive cells between the two genotypes. Quantified data are presented as mean +/- SEM of 4 animals in each group with statistics using *student's two-tailed t-test***.** ns, $p \geq 0.05$; *, $p \leq 0.05$.

Supplementary Fig. S4

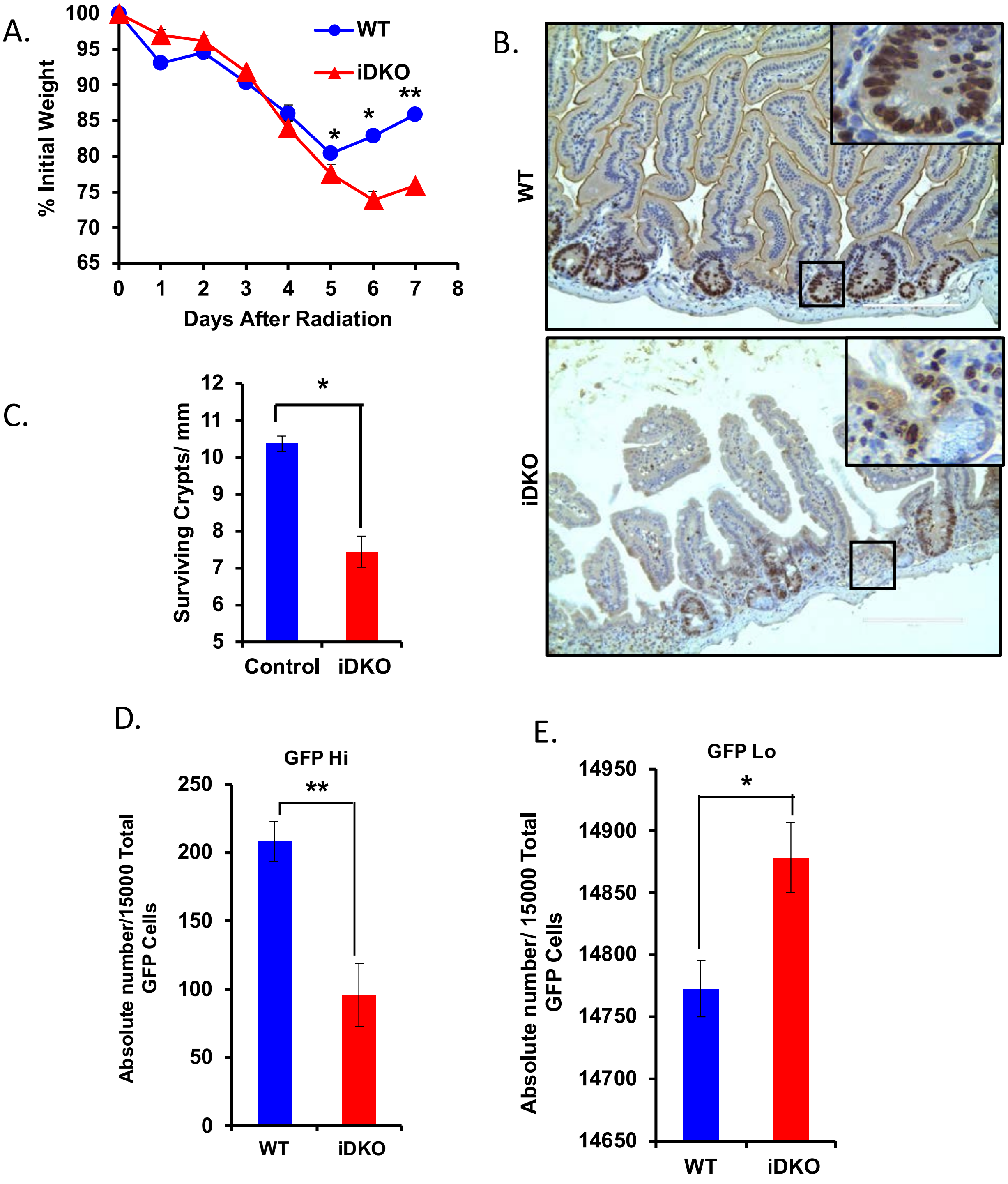

**Supplementary Figure S4. Delayed intestinal epithelial recovery from radiation injury and loss of stem cell numbers upon Lgr5+ ISC-specific *Cbl/Cblb* iDKO.** (A) Recovery from weight loss was monitored over 7 days post-radiation as a surrogate marker of recovery from abdominal radiation. *Cbl/Cblb* iDKO mice showed a significant lag in weight gain post radiation injury compared to control mice. (B, C) Ki67 staining of intestinal sections harvested on day 7 post-radiation was carried out to visualize proliferating crypts as a measure of post-radiation recovery. Representative images are shown in (B) and highlight the markedly reduced numbers of proliferating crypts in *Cbl/Cblb* iDKO mouse intestines. Quantification of proliferating crypts showed a significant reduction in *Cbl/Cblb* iDKO mice. (D, E) Intestinal epithelial cells of control and *Cbl/Cblb* iDKO mice isolated on day 5 post-TAM and radiation exposure were analyzed by FACS to quantify the absolute numbers of GFP Hi (stem) and GFP Lo (progenitor) cell populations. Quantification showed a significant drop in the GFP Hi (D) and a mild increase in the GFP Lo (E) cell numbers in *Cbl/Cblb* iDKO mice as compared to controls. Quantified data are presented as mean +/- SEM of 5 animals in each group, with statistics using *student's two-tailed t-test***.** ns, $p \geq 0.05$; *, $p \leq 0.05$; **, $p \leq 0.01$.

### Supplementary Fig. S5

A.

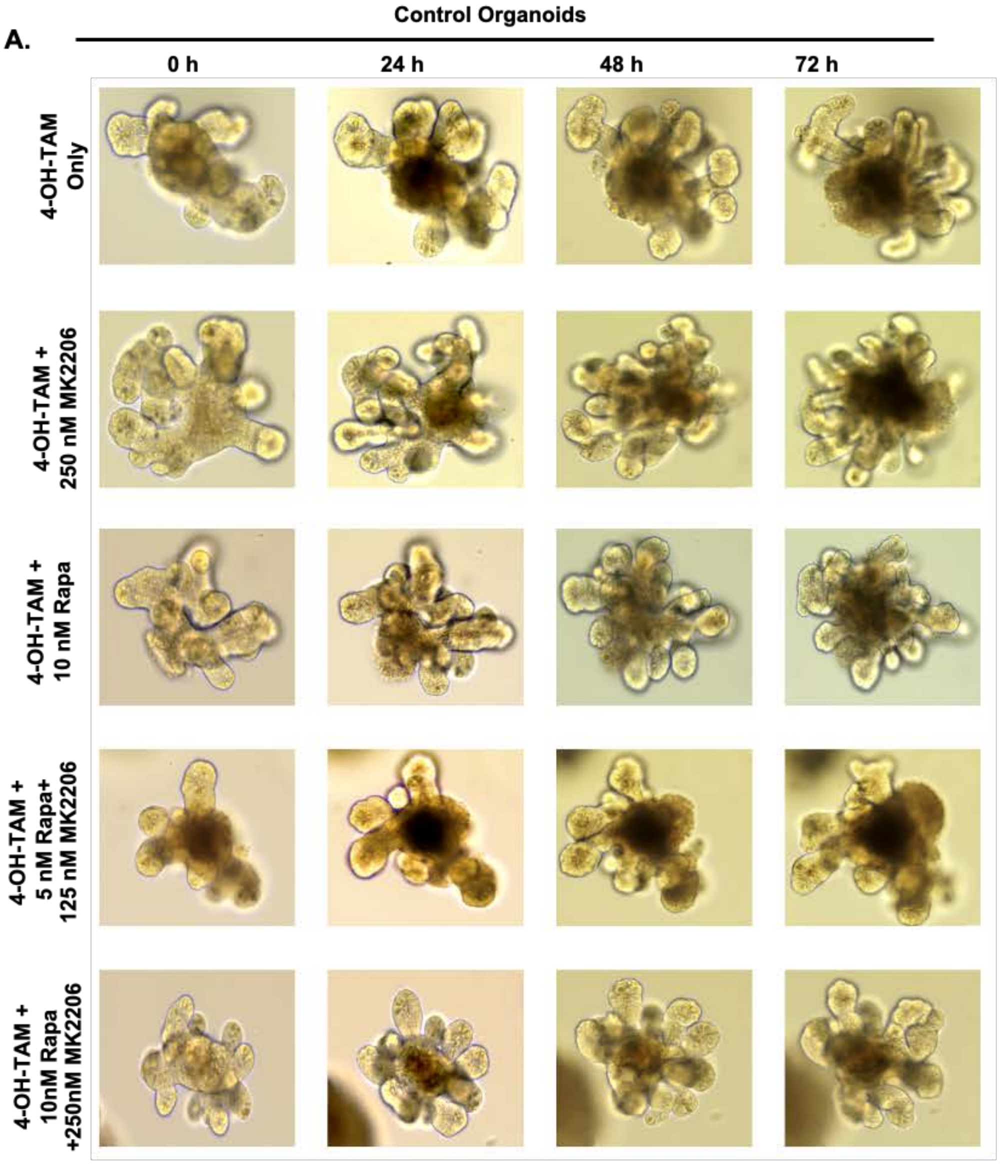


B.

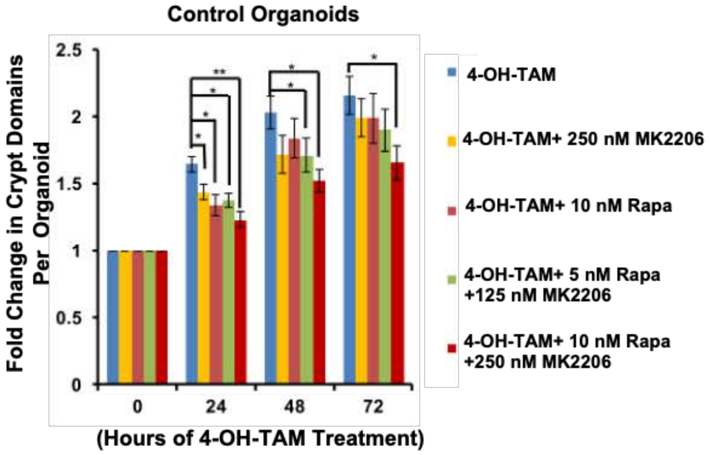

**Supplementary Figure S5. Akt/mTOR pathway inhibition does not affect the architecture of control organoids.** (A) FF Control organoids treated as part of the experiments presented in Fig. 7 to assess the effects of inhibitors on control organoid growth. Bright field images were captured every 24 h for ten different organoids per treatment condition up to 72 hours. Organoid structures were maintained under all treatment conditions with intact crypt-villus domain architecture throughout the treatment (A). Quantification of budding (crypt domains) revealed continued increase under all conditions but organoids cultured with Akt/mTOR inhibitors showed significantly reduced crypt budding as compared to 4-OH-TAM alone (B). Fold change in crypt domain quantified data of FF control organoids is presented as mean +/- SEM of 3 experiments with statistics using *student's two-tailed t-test***.** ns, $p \leq 0.05$, *; $p \leq 0.01$, ***.*
